# Thresholding of cryo-EM density maps by false discovery rate control

**DOI:** 10.1101/374546

**Authors:** Maximilian Beckers, Arjen J. Jakobi, Carsten Sachse

**Affiliations:** European Molecular Biology Laboratory (EMBL), Structural and Computational Biology Unit, Meyerhofstraße 1, 69117 Heidelberg, Germany; Candidate for Joint PhD degree from EMBL and Heidelberg University, Faculty of Biosciences; European Molecular Biology Laboratory (EMBL), Hamburg Unit c/o DESY, Notkestraße 85, 22607 Hamburg, Germany; The Hamburg Centre for Ultrafast Imaging (CUI), Luruper Chaussee 149, 22761 Hamburg, Germany; current address: Kavli Institute of Nanoscience, Department of Bionanoscience, Delft University of Technology, Van der Maasweg 9, 2629 HZ Delft, Netherlands

**Keywords:** Electron cryo-microscopy, signal detection, false discovery rate, cryo-EM density, subtomogram averaging, local resolution, ligand binding

## Abstract

Cryo-EM now commonly generates close-to-atomic resolution as well as intermediate resolution maps from macromolecules observed in isolation and in situ. Interpreting these maps remains a challenging task due to poor signal in the highest resolution shells and the necessity to select a threshold for density analysis. In order to facilitate this process, we developed a statistical framework for the generation of confidence maps by multiple hypothesis testing and false discovery rate (FDR) control. In this way, 3D confidence maps contain separated signal from background noise in the form of local detection rates of EM density values. We demonstrate that confidence maps and FDR-based thresholding can be used for the interpretation of near-atomic resolution single-particle structures as well as lower resolution maps determined by subtomogram averaging. Confidence maps represent a conservative way of interpreting molecular structures due to minimized noise. At the same time they provide a detection error with respect to background noise, which is associated with the density and particularly beneficial for the interpretation of weaker cryo-EM densities in cases of conformational flexibility and lower occupancy of bound molecules and ions to the structure.

## 1 Introduction

Cryo-EM based structure determination has undergone remarkable technological advances over the last several years leading to a sudden multiplication of near-atomic resolution structures (Patwardhan, 2017). Before these transformative changes, only highly regular specimens such as helical or icosahedral viruses were resolved at such detail (Unwin, 2005; Sachse *et al.*, 2007; Zhang *et al.*, 2008; Yonekura *et al.*, 2003; Yu *et al.*, 2008; Ge & Zhou, 2011). With the advent of direct electron detectors (McMullan *et al.*, 2016) and simultaneous improvements in image processing software (Scheres, 2012*b*; Lyumkis *et al.*, 2013; Punjani *et al.*, 2017), smaller, less regular and more heterogeneous single-particle specimens became amenable to be routinely imaged below 4 Å resolution (Bai *et al.*, 2013; Li *et al.*, 2013; Liao *et al.*, 2013). Recently, the highest resolution structures have become available at ~2 Å resolution (Merk *et al.*, 2016; Bartesaghi *et al.*, 2018; 2015) and sub-4 Å structures below 100 kDa from images obtained with and without an optical phase plate have been resolved (Merk *et al.*, 2016; Khoshouei *et al.*, 2017). These studies established the technical routines for determining atomic models of structures that were previously thought to be impossible to be resolved by cryo-EM or any other technique (Bai *et al.*, 2015; Galej *et al.*, 2016; Fitzpatrick *et al.*, 2017; Gremer *et al.*, 2017). Electron tomography is the visualization technique of choice for more complex samples including the cellular environment. Due to the poor signal-to-noise ratio (SNR) individual tomograms suffer from substantial noise artifacts. In case tomograms contain identical molecular units they can be averaged by orientationally aligning particle volumes (Briggs, 2013). Recently, with the increase of data quality and improved image processing routines, this approach also yielded near-atomic resolution maps from HIV capsid (Schur *et al.*, 2016).

The resulting reconstructions regardless of whether they originate from single-particle and subtomogram averaging are inherently limited in resolution and suffer from contrast loss at high resolution (Rosenthal & Henderson, 2003). In the raw reconstructions, the high-resolution features are barely visible as the amplitudes follow an exponential decay described by the B-factor quantity that combines effects of radiation damage, imperfect detectors, computational inaccuracies and molecular flexibility. The Fourier shell correlation (FSC) is the accepted procedure to estimate resolution (Saxton & Baumeister, 1982; van Heel *et al.*, 1982; Rosenthal & Henderson, 2003) and can be compared with a given spectral signal-to-noise ratio (SSNR) (Penczek, 2002). Consequently, B-factor compensation by sharpening is essential and common practice. Sharpening is combined with signal-to-noise weighting to limit the enhancement of noise features (Rosenthal & Henderson, 2003). Based on sharpened maps, atomic models are built and further improved by real-space or Fourier-space coordinate refinement (Adams *et al.*, 2010; Murshudov, 2016). This process is particularly challenging at resolutions between 3 and 5 Å commonly achieved in cryo-EM. Recently, we proposed a method to sharpen maps by using local radial amplitude profiles computed from refined atomic models (Jakobi *et al.*, 2017). This method facilitates interpretation of densities with resolution variation, but also requires the prior knowledge of a starting atomic model with correctly refined atomic B-factors. Despite this advance, a more general approach is needed at the initial stages of density interpretation in particular in the absence of prior model information. Tracing of amino acids derived from the primary structure as well as placing non-protein components into density maps remains a laborious and time-consuming task. In particular, the EM density contains a large dynamic range of gray values for which only a small percentage of voxels is relevant for the interpretation using isosurface-rendered thresholded representations. In practice, the process of choosing a threshold is helped by the empirical recognition of binary density features matching those of expected protein features at the given resolution. Therefore, it would be desirable to have more robust density thresholding methods at hand to reduce subjectivity and provide statistical guidance in deciding which map features are considered significant with respect to background noise.

Extracting significant information from noisy data is a common problem in many fields of science. The simplest approach is based on thresholding corresponding to multiples of a standard deviation *σ* from an expected mean value. The experimental values are only considered significant when above and rejected as noise when below this threshold. In X-ray crystallography and cryo-EM, this *σ*-approach is commonly applied to the determined maps and *σ*-thresholds are often reported when isosurface renderings of the density are displayed. In EM maps in particular, the *σ*-levels reported for interpretation are not universal and will be chosen by the interpreter as they vary from structure to structure between 1 and 5 multiples of *σ* and often to a smaller extent within the structure. The reason for the observed variation is that the high-resolution amplitudes of density peaks are very weak and can be compromised by noise after sharpening. In statistical theory, it has been recognized that the simple *σ*-method tends to increase the probability of declaring significance erroneously with larger number of tests (Miller *et al.*, 2001), which is referred to as the multiple testing problem. To account for this effect, the probability of correct detection could be increased by controlling the false discovery rate (FDR) (Benjamini & Hochberg, 1995). This statistical procedure has been applied to noisy images in astronomy (Miller *et al.*, 2001) and to time recordings of brain magnetic resonance images (Genovese *et al.*, 2002) to better discriminate signal from noise.

Due to the low SNRs of cryo-EM maps at high resolution, separating signal from noise remains a daunting task. At present, the visualization and interpretation of the density requires experience of the operator and thus relies on subjectively chosen isosurface thresholds. As sharpening procedures also amplify noise alongside the high-resolution signal, a more robust assessment of the statistical significance of those features by a particular detection error is desirable. Here, we propose to apply the statistical framework of multiple hypothesis testing by controlling the FDR to cryo-EM maps. The resulting maps we refer to as confidence maps represent the FDR of a per-voxel basis and allow the separation of signal from noise background. Confidence maps provide complimentary information to EM densities from single-particle reconstructions and subtomogram averaging as they allow detection of particularly weak features based on statistical significance measures.

## 2 Methods

### 2.1. Statistical framework

In order to overcome limitations in interpreting density features with respect to significance, we apply multiple hypothesis testing using FDR control to cryo-EM maps. In this workflow, we estimate the noise distribution from the background of a sharpened cryo-EM map, apply subsequent statistical hypothesis testing for each voxel and control the FDR (**Fig. 1a**). For the background noise, we assume a Gaussian distribution or if required an empirical density distribution where the mean and variance of the noise is estimated from four independent density cubes outside the particle density along the central *x*, *y* and *z* axes (**Fig. 1b**). Subsequently, these estimates are used to obtain upper bounds to assess signal from the particle with respect to background noise (see Appendix). In addition, we assume that cryo-EM density to be interpreted is of positive signal (see Results). Therefore, statistical hypothesis tests are carried out by one-sided testing. To account for the total number of voxels and the dependency between voxels, *p*-values are further corrected by means of a FDR control procedure according to Benjamini and Yekutieli (Benjamini & Yekutieli, 2001). The FDR-adjusted *p*-values (or *q*-values) of each voxel are directly interpretable as the maximum fraction of voxels that have been mistakenly assigned to signal over the background.

**Fig. 1.**
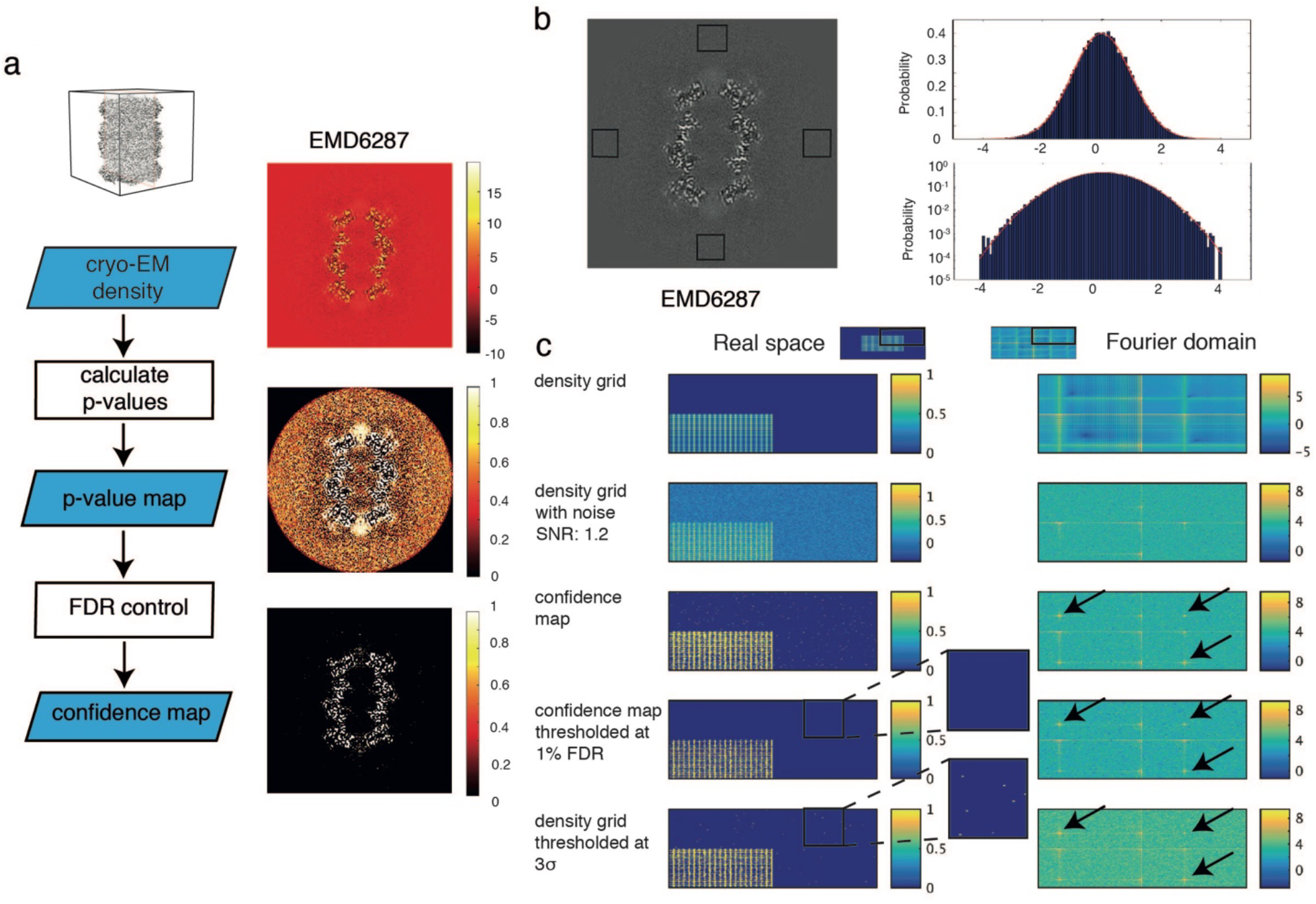
False discovery rate (FDR) analysis of cryo-EM maps. (a) Left. Flowchart of confidence map generation: the cryo-EM map is converted to *p*-values and finally FDR controlled. Right. Slice views through a cryo-EM map of 20S proteasome (EMD6287) depicted at the respective stages of the algorithm (blue boxes) on the left. Note, the strong increase in contrast when the sharpened map is converted to the confidence map. (b) Left. Estimation of the background noise from windows (red) outside the particle. Right. Histograms (top. probability in linear scale, bottom. probability in log-scale) of the background window together with the probability density function of the estimated Gaussian distribution. (c) Evaluation of the algorithm on a simulated 2D density grid. The upper right quadrant of images in real space (left column) together with the corresponding power spectrum in the Fourier domain (right column) are displayed. Density grid with added normally distributed noise at a signal-to-noise ratio of 1.2 leads to loss of contrast at high resolution. Confidence maps recapitulate these high-resolution features (arrows), showing that high-resolution signal is detected with high sensitivity. FDR thresholding at 1 % recovers a similar binary grid in comparison with 3*σ*-thresholding while minimizing noise contributions while minimizing detected noise (zoomed insets).

As *q*-values of the respective voxels provide a well-established detection measure, we further explored its use for density presentation and thresholding. Based on the FDR, we inverted the map values to the positive predictive value (PPV) by *PPV = 1 – FDR*. When the map is thresholded at PPV of 0.99, at least 99% of the binarized voxels are truly positive density signal within the map, corresponding to a FDR of 1%. We term these maps confidence maps, referring to the fact that PPVs serve as a measure of the confidence by which we can discriminate signal from the noise. These confidence maps can then be visualized like usual cryo-EM maps with common visualization software, with the difference that the threshold for visualization is now given by 1–FDR rather than the density potential.

### 2.2. Simulations

The simulated images were 400×400 pixels in size. The scaled grid was generated by adding two orthogonal two-dimensional cosine waves with a period of 5 pixels, where all values smaller than zero were set to 0, and multiplying the sum by a factor of 0.5 in order to scale the maximum to 1. The scaled grid was 200×200 in size and embedded in the center of the 400×400 image. Gaussian distributed noise with a mean of 0 and given variance of 0.01 (**Fig. 1c**), 0.1 (**Fig. S1a**) or 1.33 (**Fig. S1b**), respectively, was added to the grid image. Mean and variance for the multiple testing procedure were estimated outside the scaled grid and the FDR-procedure was carried out as described. Simulations were implemented in MATLAB (Mathworks Inc.).

### 2.3. Software

The algorithm is implemented in Python, based on NumPy (Walt *et al.*, 2011) and the mrcfile I/O library (Burnley *et al.*, 2017). Local resolutions were calculated using ResMap (Kucukelbir *et al.*, 2014). The software is available at https://git.embl.de/mbeckers/FDRthresholding. Figures were prepared with UCSF Chimera (Pettersen *et al.*, 2004).

## 3 Results

### 3.1. FDR-based hypothesis testing yields improved signal detection in simulations

In order to evaluate the principal performance of the proposed method on simulated data, we prepared a two-dimensional grid of continuous density waves (**Fig. 1c, left**). We added white noise to a series of test images containing SNRs between 3.9 and 0.3 as they occur in high-resolution shells of 3D reconstructions when the FSC curve drops from 0.67 up to 0.143 often reported as the resolution cutoff. First, we generated a test image with a SNR of 1.2 and noted that in the power spectrum computed from the simulated noise images, signal from high-resolution features cannot be detected although being present in the noise-free power spectrum (**Fig. 1c, right**). The detection of these high-resolution features, however, can be recovered from the corresponding confidence images that we generated as described above, even at SNRs ranging between 3.9 and 0.3 (**Fig. S1ab**). When comparing images thresholded at conventional 3.0*σ* levels with confidence images thresholded at a PPV of 0.99 or FDR of 0.01 (from here on referred to as 1% FDR), we note that FDR-controlled thresholding allows more faithful detection of weak density features closer to noise levels. In this way, the density transformation to confidence images minimizes false positive detection of pixels and improves the peak precision as adjacent noise peaks are suppressed (**Fig. S2**).

### 3.2. Choice of positive density model with Gaussian background noise

Although the model of Gaussian noise is often used to approximate background noise in cryo-EM images and maps (Sigworth, 1998; Scheres, 2012*a*; Kucukelbir *et al.*, 2014; Vilas *et al.*, 2018), it is important to analyze actual maps to better understand deviations from this assumption. For this purpose, we analyzed a total of 32 deposited cryo-EM densities from 2 to 8 Å resolution and compared the empirical cumulative density function (CDF) with the ideal Gaussian CDF (**Fig. S3a**). It is apparent that all of them follow the ideal Gaussian CDF closely. For each map, we assessed normality by Anderson-Darling hypothesis testing (Anderson & Darling, 1954) and found that 75% and 87.5 % of the maps do not significantly deviate from normality when conservative thresholds corresponding of 1% and 0.1% Family Wise Error Rates (FWER) are chosen (**Fig. S3b**). One of the reasons for the observed deviations from an idealized Gaussian distribution is a result of the 3D reconstruction procedure. In principle, when truly aligned images containing white Gaussian noise are combined by linear inversion, the obtained 3D volume will also have Gaussian distribution. In practice, in cases when uncertainties reside on the 5 orientation parameters, background noise is not necessarily Gaussian distributed. Moreover, resulting 3D reconstructions will contain local correlations, i.e. “colored noise”. Therefore, we analyzed the resulting noise of 3D reconstructions generated from pure noise images with even angular sampling. The resulting amplitude spectrum shows that it differs from pure white noise due to correlations between adjacent pixels (**Fig. S3c, left**). Furthermore, variances estimated for each voxel from 900 reconstructions show that they can be approximated uniform over the central sphere (**Fig. S3c, right**).

For the map of EMD-6287, which according to the Anderson-Darling test deviates strongly from normality, we generated a confidence map using the Gaussian and the empirical CDF. We inspected these confidence maps (**Fig. S3d**) and find that the visual agreement between the two maps is very high. To highlight potential differences, we computed a difference map between the two confidence maps created by the two approaches and observe no systematic variation when deviation from normality is assumed. Therefore, for interpreting confidence maps, small deviations from normality do not appear to have practical limitations. In order to rule out any potential unforeseen effects when maps deviate more strongly, we routinely implemented the monitoring the degree of deviation from ideal Gaussian CDF. For instance when the deviation of the empirical CDF from the Gaussian CDF exceeds 0.01, referring to the fact that *p*-values deviate by more than 1 %, can optionally use the empirical CDF for the generation of confidence maps.

The second assumption of the proposed confidence map assumes that protein gives rise to positive density in cryo-EM maps. When inspecting EM density maps, it is evident that not all signal present in the map is positive. Therefore, we analyzed whether significant negative densities can be detected in confidence maps generated from inverted densities. Indeed the confidence maps from negative densities reveal significant signal in regions between protein density often in a spatially complementary way (**Fig. S4a left**). Using the independently determined X-ray structure of the 20S proteasome (PDB code 1PMA), we tested whether negative density coincides with the atomic model. Overall, negative density has only a very small 2.5 % overlap with atoms, which is close to the predicted false discovery rate of 1 % (**Fig. S4b**). When using positive density, however, we find that a large fraction of 60 % of the PDB atoms are found in the 1 % FDR contoured confidence map and 10% of that volume is occupied by modelled atoms. In conclusion, we show that negative density presents significant signal in cryo-EM maps, however, that only a very small fraction is occupied by atoms. The largest fraction of negative densities are found next to positive protein density most likely due to the fact that the molecular density is lower than in the particle surrounding solvent area. Based on this analysis and our objective to identify those voxels that arise from protein density, we include the restraint of testing for positive signal into the generation of confidence maps and include an additional option to test for negative signal.

### 3.3. Confidence maps from near-atomic resolution maps separate signal from background suited for molecular structure interpretation

In order to assess the potential of confidence maps for the interpretation of cryo-EM densities, we applied the algorithm to the near-atomic resolution map of TMV determined at a resolution of 3.35 Å (EMD 2842) (Fromm *et al.*, 2015). Variances could be estimated reliably outside the helical rod from a range of different window sizes from 10 to 30 voxels using the cryo-EM density (**Fig. S5**). To generate the confidence map, we transformed the cryo-EM density to *p*-values and subsequently to confidence maps in an equivalent manner to the simulated confidence images above. Next, we inspected a longitudinal TMV section through the four helical bundle of the coat protein and compared the confidence map with the cryo-EM density (**Fig. 2a and b**). The confidence map revealed backbone traces that contain values close to 1 corresponding to the helical pitch of the LR helix. They clearly stand out with respect to background noise that is suppressed towards values of 0. The associated histogram of the confidence map revealed a strong peak beyond 0.99 PPV or below 1 % FDR separating signal over background and thresholding 5.7 % of voxels within the density. In the case of the deposited cryo-EM map, the subjectively fine-tuned and recommended 1.2 *σ* threshold also yielded a recognizable outline of helical pitch contours while detecting only 3.7 % of voxels from the density. In analogy to isosurface-rendered cryo-EM densities, confidence map exhibit recognizable structural details, such as the α-helical pitch and many side chains of the central helices (**Fig. 2c**). When applying a lower FDR of 0.01 %, polypeptide density becomes discontinuous and smaller density features disappear. When going to higher FDR thresholds such as 10 %, noise starts to be included in the density. At the recommended 1 % FDR threshold, the appearance of noise is minimal and well controlled in confidence maps. This is in contrast to cryo-EM densities, where the appearance of noise is very sensitive to small changes in threshold level in particular at lower *σ*. In fact, the recommended 1.2 *σ* contour includes only 52 % of the atoms of the model whereas the 1 % FDR threshold contour already contains 73 % with minimized noise. In order to include the same amount of atoms in a contour, a threshold of 0.7 *σ* would be required, which at the same time will lead to a noticeable increase of obstructing noise. Furthermore, we also examined two additional confidence maps from EMDB model challenge targets determined at near-atomic resolution: 20S proteasome (Campbell *et al.*, 2015) and γ-secretase (Bai *et al.*, 2015) (**Fig. S6a and S6b**). These confidence maps confirm the previous observation that when displayed at FDR levels of 1 %, they provide structural details at near-atomic resolution while effectively separating signal from noise.

**Fig. 2.**
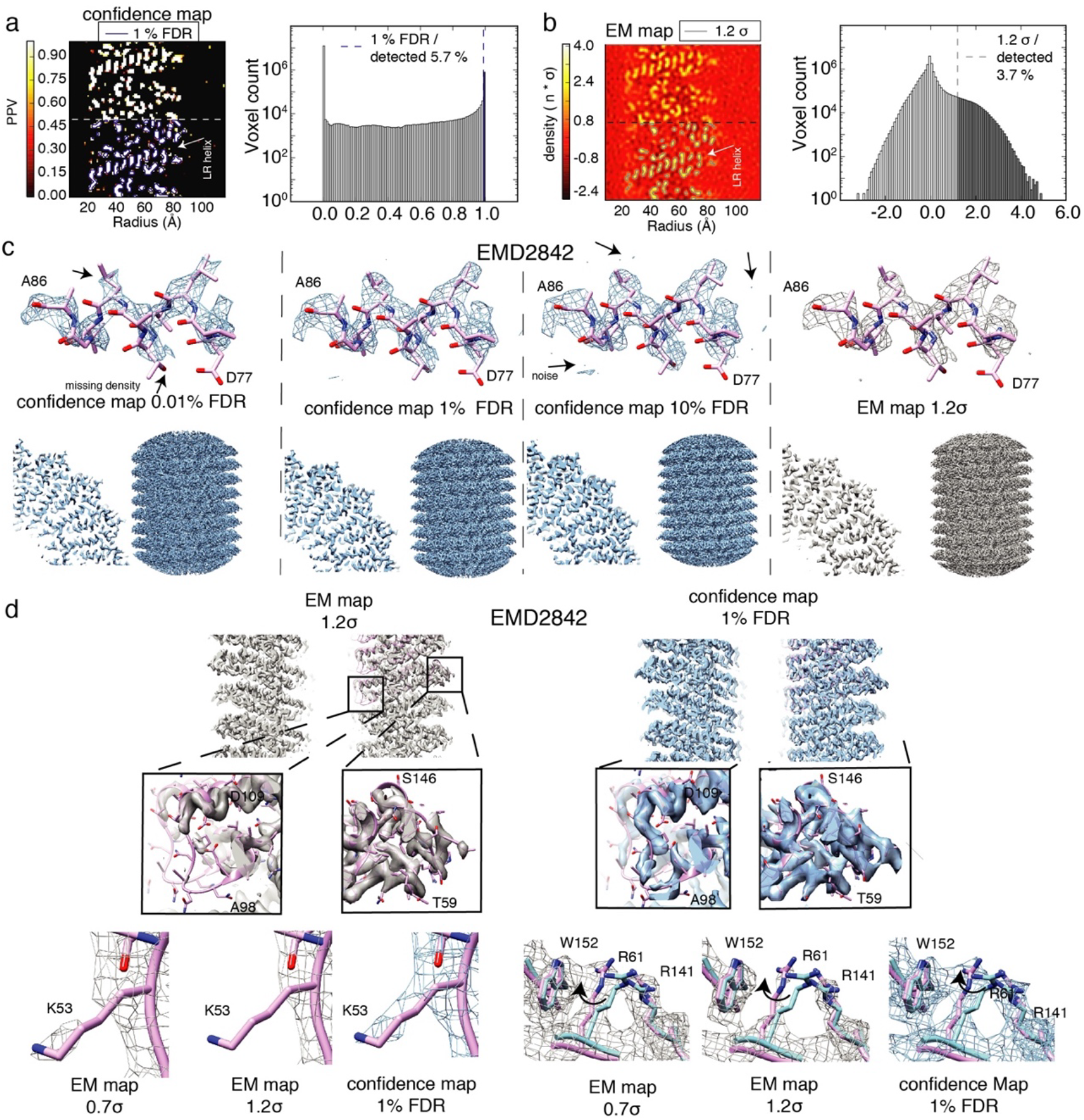
Confidence maps separate signal from noise for molecular density interpretation. (a) Left. Confidence map with longitudinal section through TMV coat protein displayed indicating α-helical pitch of LR helix. Lower half shows the chosen contour at 1 % FDR in blue with 5.7 % of voxels detected. Right. Corresponding histogram of confidence map with signal separated above 0.99 PPV (1 % FDR). (b) Left. Same section as in (a) from cryo-EM density and the recommended threshold contoured at 1.2 *σ* in gray with 3.7 % of voxels detected. Right. Corresponding histogram of cryo-EM density with thresholded values displayed in gray. (c) Isosurface rendered of thresholded confidence maps at 0.01 %, 1 % and 10 % FDR (left, center left, center right) shown in blue and sharpened cryo-EM density with 1.2*σ* threshold (right) in gray from TMV (EMD2842). Shown are helix A86 – D77 (top), quarter cross section (bottom left) and side view (bottom right) of TMV map. (d) Detailed analysis of TMV density. Slice view through TMV rod with zoomed inset for inner and outer radii density (top). K53 side chain density (left) and molecular environment of R61 side chains (right) at 0.7, 1.2 *σ* threshold and at 1 % FDR confidence map.

### 3.4. Confidence maps provide a map detection error with respect to background noise

When confidence maps are generated from cryo-EM densities, the main aim of the approach is to determine a voxel-based confidence measure of molecular density signal with respect to background noise. In principle, the confidence measure could also be interpreted as a broader error estimate of the EM map referring to the rate of falsely discovered of voxels. The error, however, as it arises from a cryo-EM experiment is a comprehensive quantity, which results from multiple contributions in the form of the solvent scattering, detector noise as well as computational sources of alignment and reconstruction algorithms in addition to variation of signal by multiple molecular conformations and radiation damage effects (Frank & Liu, 1995; Penczek *et al.*, 2006). Estimating the complete series of error contributions including signal variation is currently not possible in the context of common cryo-EM collection schemes. Therefore, the most straightforward way of estimating noise is measuring the variance of the map solvent area. This variance mainly captures errors as they arise from detector noise and solvent scattering while neglecting contributions of computation and local molecular variations. Detector noise can be considered to be distributed uniformly over the 3D reconstruction whereas solvent scattering distribution will not be uniform as pure solvent noise next to the particle is higher when compared with solvent noise projected through the particle due to solvent displacement and variations of water thickness in the particle view (Penczek, 2010). Consequently, measuring noise in the solvent area of cryo-EM maps, will lead to an effective overestimation of noise and therefore to an underestimation of confidence (see Appendix Proposition 1). Although these deviations from a uniform Gaussian noise model do not allow absolute error determination, in practice estimating solvent variance can be used as conservative upper bounds for error rates without including errors arising from computation and molecular variation. In conclusion, the error as it arises from confidence maps should be considered a map detection error with respect to background noise that can assist in the interpretation of cryo-EM densities.

### 3.5. Robustness of FDR-controlled density transformation

In order to test the robustness of the approach, we systematically assessed the effects of the required input on the resulting confidence map. First, we tested the influence of severely underestimating noise for confidence map generation by using the 1/2 or 3/4 of the determined variance of the 20S proteasome densities (**Fig. S6c**). The resulting confidence maps displayed at 1 % FDR revealed excessive declaration of background as signal, which poses a principal risk for over-interpretation. This principal risk, however, is less relevant, when the here proposed variance measurements outside the particle is used as we tend to overestimate noise (see above and Appendix Proposition 1). Therefore, we tested the effect of overestimating the variance by 1.25, 2 and 8 fold and generated confidence maps according to the defined procedure. The resulting confidence maps show the disappearance of map features at the 1 % FDR threshold only when the variance is severely overestimated by a factor of 8 but for small overestimations is hardly noticeable in the map appearance. Another important noise-related parameter prior to the proposed procedure is the applied sharpening level. Therefore, we tested a series of B-factors from 0 to −250 Å^2^ applied to the 20S proteasome maps and converted them into confidence maps. First, with increasing negative B-factors the corresponding confidence maps displayed at 1 % FDR show loss of features due to the drop in relative significance. This is in contrast to cryo-EM densities that become severely over-sharpened and density features are dominated by noise (**Fig. S6d**). Second, when under-sharpened maps are used for noise estimation, maps will contain only low-resolution features lacking high-resolution detail at the respective significance level in analogy to cryo-EM densities. Therefore, when over-sharpened maps are used for noise estimation, confidence maps inherently avoid enhancement of noise features that could be mistakenly interpreted as signal. Although noise estimation is important for the procedure, tests show that smaller variance overestimation does not have a noticeable effect on map interpretation of 1 % FDR confidence maps. In conclusion, confidence maps represent a conservative way of displaying maps at defined significance while avoiding the problem of over-sharpening, which represents a principal benefit over visualization of *σ*-thresholded sharpened EM densities.

### 3.6. Confidence maps facilitate detection of weak density features

In order to evaluate further molecular details of the confidence map, we inspected more ambiguous density features of the TMV map. Peripheral density at lower and higher radius of the virus was notoriously difficult to interpret in previous works (Fromm *et al.*, 2015; Sachse *et al.*, 2007; Namba & Stubbs, 1986). For these regions, we found that there are well-defined features present in the 1% FDR confidence maps. Densities of the coat protein for loops Q97 – T103 located at the inner radius and T153 – G155 at the outer radius are not present in the respective EM map, but clearly traceable in the 1% FDR confidence map (**Fig. 2d, center**). In addition, side-chain density for K53 contacting the adjacent subunit was found to be clearly significant while being discontinuous in the original map (**Fig. 2d, bottom left**). Based on confidence maps, readjustment of side-chain rotamers was possible, illustrated for example by significant density for R61, which suggests a realignment of R61 to form stabilizing interactions with aromatic W152 (**Fig. 2d, bottom right**). The presented examples of TMV illustrate that confidence maps represent an alternative for density display, which can help in the process of molecular feature detection. Although threshold adjustments in cryo-EM maps can also help model interpretation in ambiguous regions and enhance weak density features, they also amplify noise features and increase the risk of noise fitting.

We also tested cases of more heterogeneous densities such as the V-ATPase SidK complex (EMD-8724), which was determined at 6.8 resolution (Zhao *et al.*, 2017). First, the deposited EM map contains very weak density of the bacterial effector SidK EM density due to low occupancy and flexible motion. The corresponding confidence map of the V-ATPase SidK complex reveals that the SidK density is not significant as continuous density when thresholded at 1% FDR as it is too noisy for further analysis (**Fig. S7a**). Below in section 3.8, we will deal with cases of local resolution and SNR variation that can be accommodated by a locally adjusted FDR procedure. Second, we analyzed confidence maps from three conformational states generated by 3D classification (EMD-8724, EMD-8725, EMD-8726). The generated confidence maps thresholded at 1 % FDR of state 1, 2 and 3 confirm previous observations using EM maps (**Fig. S7b**). Taken together, confidence maps provide an inherent significance level associated with the density and minimize false positive noise detection. In this way, confidence maps can guide atomic model interpretation of cryo-EM density maps in particular in density regions of ambiguous quality.

### 3.7. Confidence maps from subtomogram averages

We further explored whether structures determined at lower resolution may also benefit from the approach. For this purpose, we examined the *in situ* determined sub-tomogram average of the HeLa nuclear pore complex computed from 8 pore particles at 90 Å resolution (Mahamid *et al.*, 2016). The deposited map clearly shows continuous densities for the cytoplasmatic and inner ring molecules whereas density below and above the pore is noisy when visualized at a threshold of 2.0 *σ* (**Fig. 3a**). The corresponding the 1% FDR confidence map shows continuous features of the ring structure with minimized noise, which makes interpretation straightforward. In order to generate a confidence map for a subtomogram average structure, care must be taken in identifying areas of noise devoid of any signal in order to estimate the noise variance reliably (**Fig. S8a**). The same tomograms recorded from lamella of HeLa cells also yielded a subtomogram average of ER-associated ribosomes. The ribosome structure itself could be determined at 35 Å at the membrane with weak density below the membrane ascribed to a translocon-associated protein complex and an oligosaccharyltransferase (Mahamid *et al.*, 2016). The corresponding densities can only be visualized at low thresholds corresponding to 0.8 *σ* while increasing the amount of background noise and hampering molecular interpretation (**Fig. 3b**). The 1 % FDR confidence maps, however, display the additional protein complexes in the absence of noise. In this case, the confidence map discriminates between specific association of the TRAP complex and the looser association of ribosomes within the polysome assembly. Further, we examined deposited and confidence maps of the 23 Å resolution nuclear pore structure determined by subtomogram averaging (Appen *et al.*, 2015) (**Fig. 3c**). While the overall densities look very similar, we focused our comparison on ambiguous density assignment of the linker region of Nup133. Presence of density in the 1% FDR confidence maps confirms the continuity of this density stretch and the author’s interpretation of placing the Nup133 linker region connecting the N-terminal β-propeller and C-terminal α-helical domain (**Fig. 3c, upper right**). In addition, we identified additional densities in the connecting densities between the inner and nuclear as well as the inner and the cytoplasmic ring (**Fig. 3c, bottom**). Both densities are not visible at the recommended σ-threshold of 2.1 but they are reliably displayed in the 1% FDR confidence map. Taken together, confidence maps generated from lower resolution subtomogram averages as well as from near-atomic resolution reconstructions assist in the density interpretation by separating signal with respect to background noise.

**Fig. 3.**
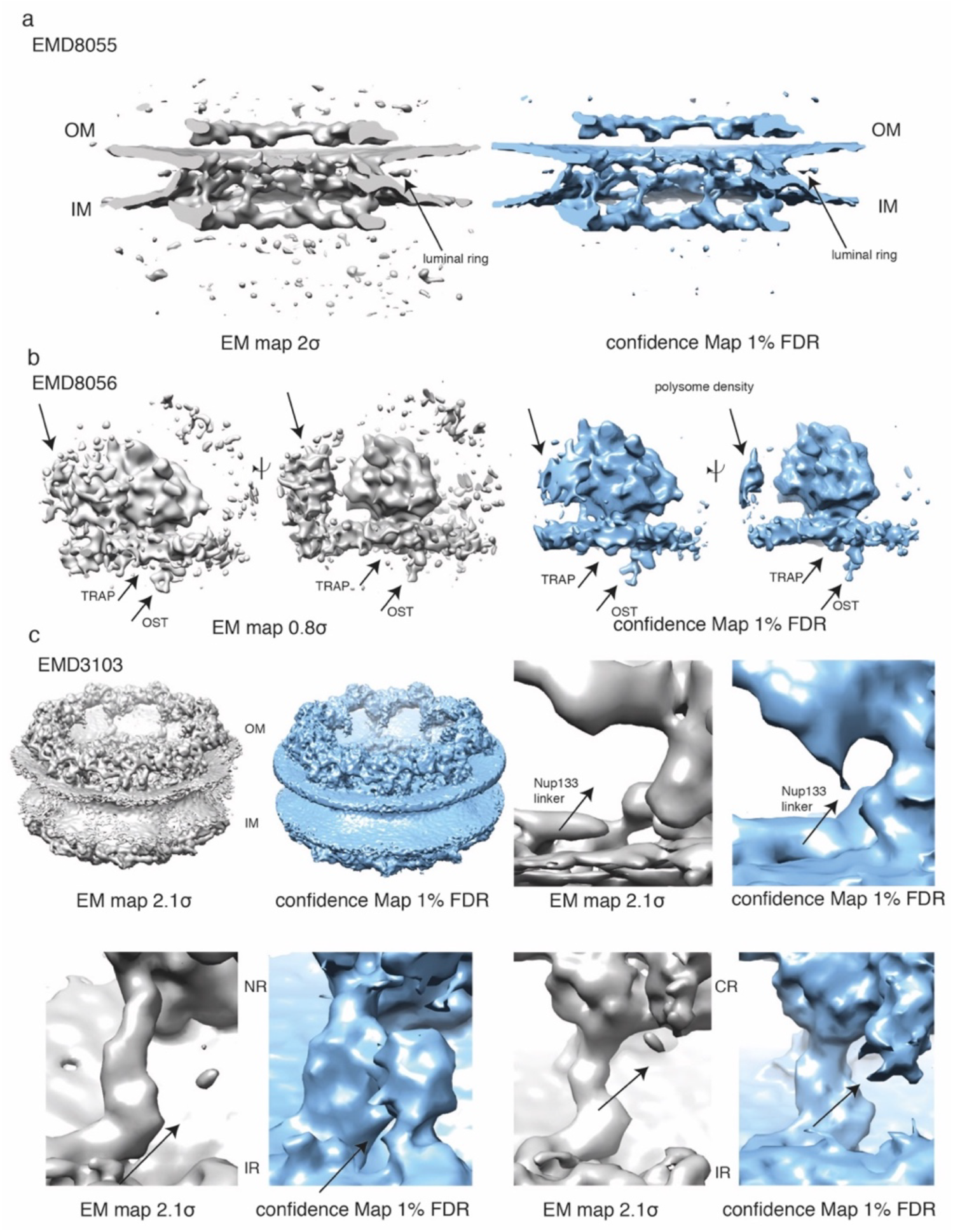
Confidence maps from subtomogram averages. (a) Nuclear pore structure at 90 Å (EMD 8055) from 8 pore particles: cryo-EM map at 2.0*σ* threshold (left, gray) and confidence map at 1 % FDR threshold (right, blue). Note, the confidence map minimizes appearance of noise. (b) ER-associated ribosome structure at 35 Å resolution (EMD 8056) in two side views at 0.8*σ* threshold (left) and 1 % FDR confidence map (right). Note, in confidence maps weaker densities assigned to peripheral protein complexes TRAP and OST (arrows) can be easily visualized in the absence of noise. (c) Nuclear pore structure at 23 Å resolution (EMD 3103) comparing cryo-EM map at 2.1*σ* threshold (left) and 1 % FDR confidence map (right). Comparison of map pairs for Nup133 linker density (top right), densities located between inner and nuclear ring (bottom left) and inner and cytoplasmic ring (bottom right). In contrast to sharpened cryo-EM maps at 2.1*σ* threshold, confidence maps show consistently densities at the connections between the inner and outer rings at 1 % FDR threshold (arrows).

### 3.8. Confidence maps benefit from local SNR adjustment in cases of resolution variation

After establishing the usefulness for maps covering a range of resolutions, we wanted to further explore how FDR-controlled confidence maps cope with large resolution differences within a single map. For this purpose, we analyzed the very high-resolution 2.2 Å map of β-galactosidase (β-gal) (EMD2984) (Bartesaghi *et al.*, 2015) in more detail as it covers resolution ranges from 2.1 to 3.8 Å. In order to reveal high-resolution details in the center of the map, high sharpening levels were required and consequently less well resolved parts in the periphery of the map resulted in over-sharpened densities. When we applied our method to the cryo-EM density volume, we found the 1% FDR confidence to be well defined in the center of the map but fading out for large parts of the periphery in support of the B-factor test series on the 20S proteasome (**Fig. S8c**). We reasoned when resolution differs across the map as a consequence of molecular flexibility and computational errors, the SNR will vary in correspondence. To compensate for these effects, noise levels can be adjusted in cryo-EM maps by applying local low-pass filtrations in Fourier space according to local resolutions (Cardone *et al.*, 2013). Consequently, a local variance can be estimated for each voxel by applying the same low-pass filter to the background noise windows (**Fig. S9a**). Application of this procedure followed by the FDR control yield a more evenly distributed 1% FDR confidence map including the β-gal periphery (**Fig. 4a, b top**). At the same time, side chain details such as holes in aromatic rings can be resolved at the same significance level as exemplified for W585 in analogy to the appropriately filtered density (**Fig. 4a, b bottom**). Closer inspection of the cryo-EM density shows that we did not observe density for the peripheral loops of the β-gal complex at the 4.5 *σ*-threshold but clearly detected continuous loop density at a FDR of 1% of the resolution-compensated confidence map (**Fig. 4c, left and right**). These observations show that the statistical power of the procedure can be improved, i.e. the amount of missed signal can be reduced in cases non-uniform noise levels, while still controlling the FDR by incorporation of local resolution information (see Appendix for detailed discussion).

**Fig. 4.**
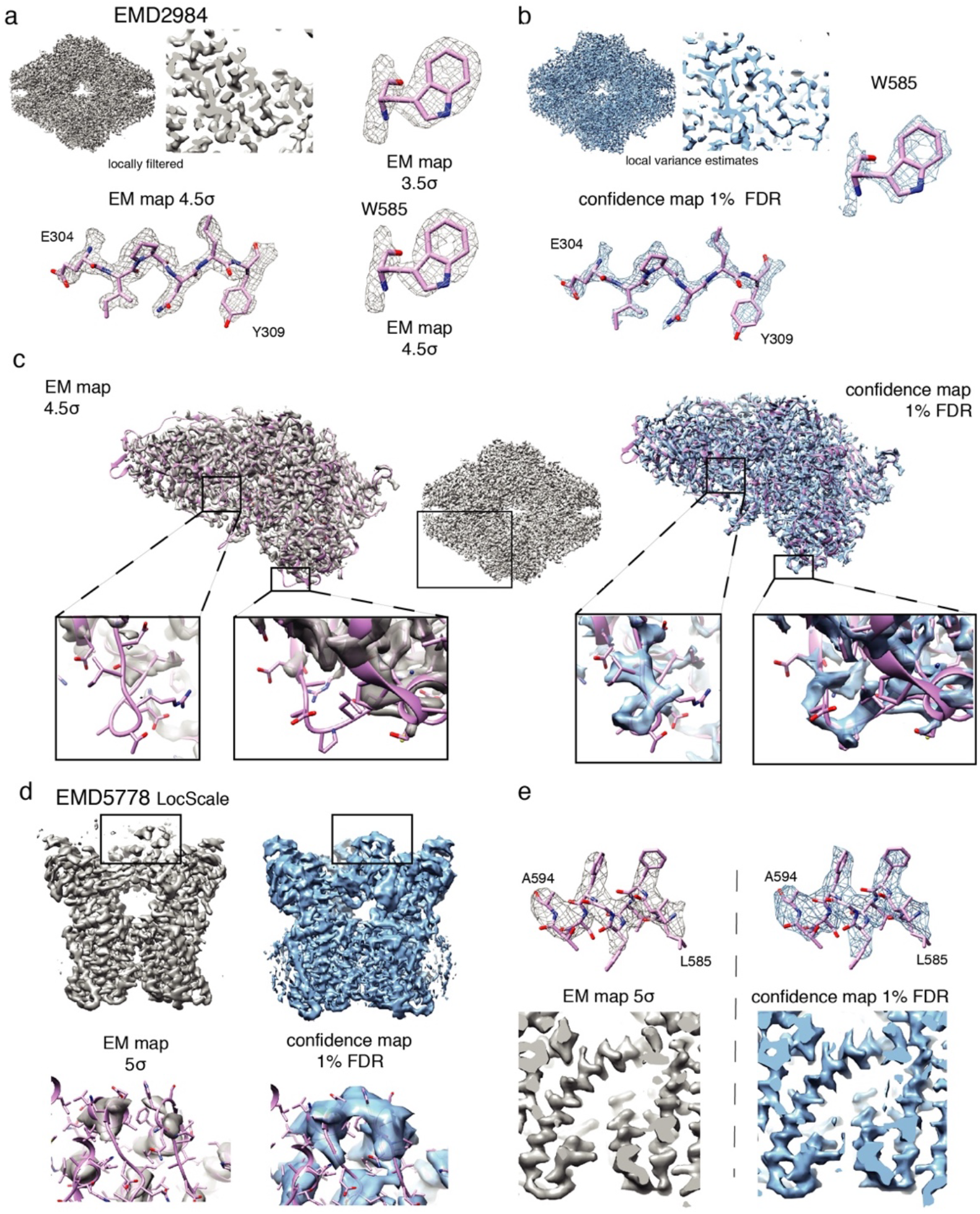
Confidence maps benefit from local SNR adjustment based on local resolution. (a) β-galactosidase (EMD 2984) locally filtered cryo-EM map left (gray) displayed at 4.5*σ* threshold and (b) confidence map (blue) including signal-to-noise adjustment based on local resolution at 1% FDR threshold (right) in side view and cross section. High resolution features like E304 – E398 and holes in aromatic rings W585 in the 3.5/4.5 *σ*-thresholded cryo-EM map (a) in comparison with the 1% FDR confidence map (b). (c) Comparison of density features from peripheral loop regions not covered by density in the locally filtered cryo-EM map (left) compared with the 1% FDR confidence map that shows densities for the respective loops. (d) TRPV1 (EMD5778) side view (top) with zoom-in to peripheral cytoplasmic domain density (bottom) comparing LocScale density displayed at 5*σ* threshold (left) and 1 %FDR confidence map. (e) Detailed density stretch A594 – L585 (top) and transmembrane helix S5 including S4-S5 linker (bottom) comparing LocScale density and 1 % FDR confidence map.

We recently introduced a local map sharpening tool for cryo-EM maps based on refined atomic B-factors (Jakobi *et al.*, 2017). When refined atomic coordinates are available, the concept of resolution-compensated confidence maps based on adjusted variances derived from local resolution filtering can be easily extended by scaling the radial amplitude falloff of the noise window against the local reference model for estimating the resulting local noise levels (**Fig. S9b**). In order to directly compare confidence maps generated by different filtering or scaling approaches, we focused on the inspection of the peripheral regions of the β-gal enzyme as the densities are weak in particular for loops extending from the particle. When we compared the confidence map of this region generated using the local resolution filtering with the original confidence map, we confirm the observation that adjustments according to local resolutions improve the density connectivity (**Fig. S10a, b**). When we used the local amplitude scaling approach, we obtained a confidence map with improved density coverage when compared with the original confidence map but less coverage when using local resolution filtering (**Fig. S10b, c**). In combination, when local variance is estimated based on local amplitude scaling and filtering, we find optimal coverage of density and the atomic model (**Fig. S10d**). Another example from the EMDB model challenge is the TRPV1 channel determined at 3.4 Å resolution (EMD5578) (Liao *et al.*, 2013). The structure contains a well-defined transmembrane region and a more flexible cytoplasmic domain that is less well resolved. The application of locally adjusted SNRs to the confidence map yields a map with well interpretable density including molecular details (**Fig. 4d and 4e**). In analogy to the examples above, the cytoplasmic domain is only visible at lower thresholds than the core of the protein. The 1 % FDR confidence map captures all density occupied by the protein including the more flexible regions at the cytoplasmic domain. The example of the TRPV1 channel confirms the observation of β-gal that local resolution differences need to be taken into account for correct generation of confidence maps. When maps exhibit strong local variation of noise due to molecular flexibility and computational errors, local variances can be estimated based on local resolution measurements or on local sharpening procedures and yield well-interpretable confidence maps at a single FDR threshold.

### 3.9. Confidence maps confirm detection of bound molecules

The majority of near-atomic resolution maps obtained by cryo-EM are in the resolution range between 3 and 4.5 Å. Although main-chain and large side-chain density can often be modeled reliably, smaller side chains and ordered non-protein components such as water molecules and ions are inherently difficult to model at these resolutions and pose the risk of noise fitting. Therefore, we investigated whether confidence maps can help to mitigate this problem and inspected a putative Mg^2+^ site coordinated by E416, E461, H418 and three additional H_2_O molecules inside of the β-gal enzyme. We rigidly placed of the Mg^2+^ ion and coordinated water molecules based on the 1.6 Å resolution X-ray crystal structure (Wheatley *et al.*, 2015) (PDB 4ttg) and superposed them onto the deposited EM density map. The map at lower 3.5 *σ* threshold shows convincing density for only two out three water molecules. (**Fig. 5a top left**). In contrast, the 1% FDR confidence map based on local variance estimation reveals distinct density peaks for all three suspected H_2_O molecules (**Fig. 5a top right**). Furthermore, β-gal had been imaged in the presence of the small molecule inhibitor PETG. Locating and conformational modeling of the ligand remains challenging due to flexibility and lower occupancy (**Fig. 5a bottom left**). Ligand placement is facilitated using confidence maps, with density well resolved for the complete small molecule inhibitor (**Fig. 5a bottom right**). The confidence density confirms previous re-refinement of the inhibitor position and conformation (Jakobi *et al.*, 2017). In addition, we also tested whether detection of smaller ions can be facilitated by confidence maps. For this purpose, we turned again to the TRPV1 channel and inspected the density surrounding G643 known as the selectivity filter for the ions passing the channel. The deposited map reveals a density peak in the symmetry center that is compatible with a small ion. In support, the confidence map also shows a density peak at the same position supporting the presence of an ion with a confidence of 1 % FDR (**Fig. 5b bottom right**). In correspondence, there are multiple cryo-EM structures reporting putative ion densities along an array of carbonyl forming an inner cavity of the channel (Lee & MacKinnon, 2017; McGoldrick *et al.*, 2018). Closer inspection of the γ-secretase complex reveals significant density for a membrane-embedded phosphatidylcholine (PC) lipid molecule. In order to detect the two PC acyl chains, the deposited EM map requires thresholding at two different *σ*-levels of 4 and 5 presumably due to differences in chain mobility (**Fig. 5c**). In contrast, the corresponding 1% FDR confidence map encompasses most of the density of two acyl chains without the need of threshold adjustments. In conclusion, confidence maps from cryo-EM structures possess minimized noise and can be directly used to evaluate the significance of density features to be present by providing a map detection error that e.g. 1 % of the peaks are expected to be falsely discovered. Using complementary information for the interpretation of cryo-EM structures will help to reduce subjectivity involved in the process of density interpretation.

**Fig. 5.**
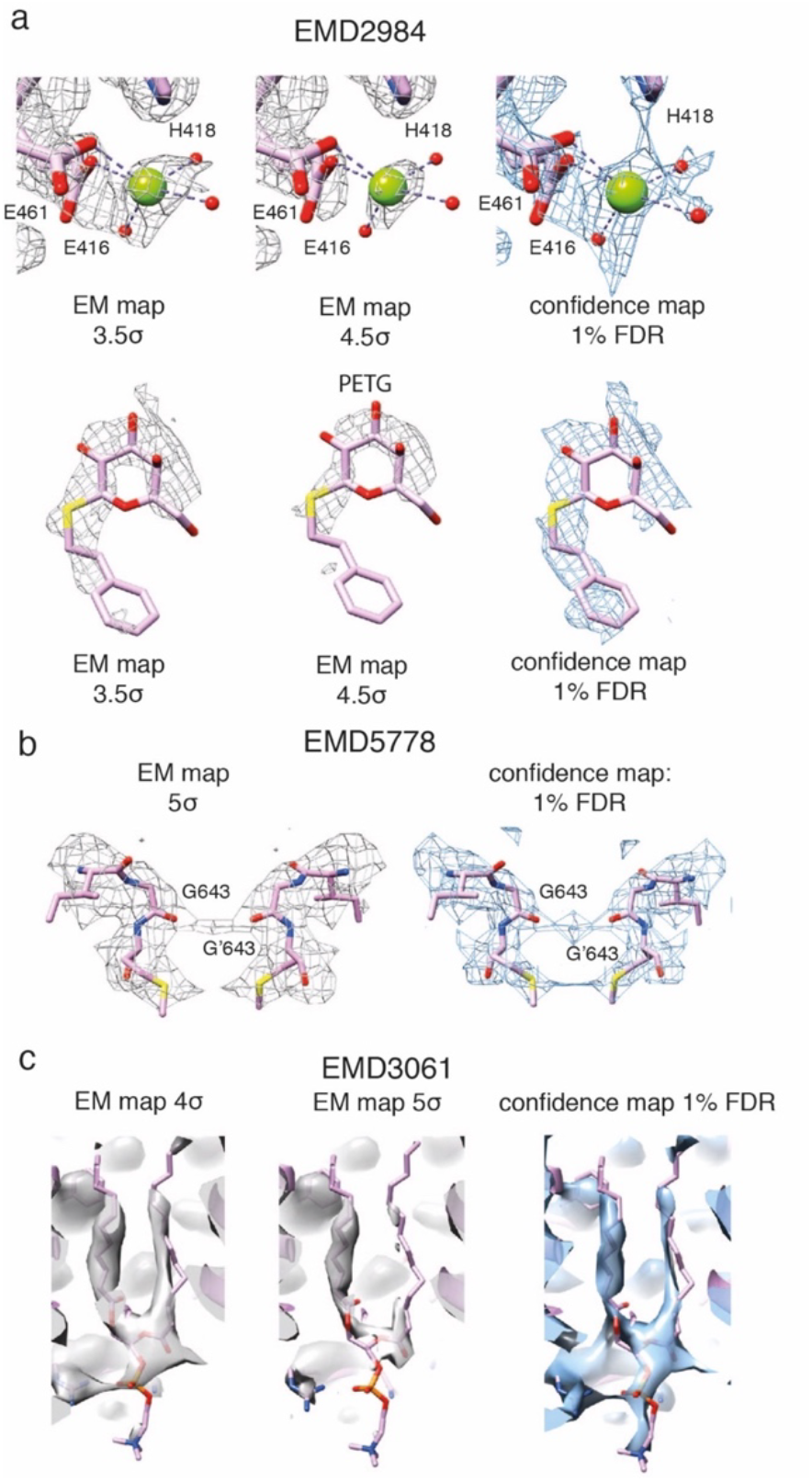
Confidence maps confirm localization of non-protein components. (a) β-galactosidase (EMD 2984) with 3.5/4.5 *σ*-thresholded cryo-EM map (left, center, gray) and 1 % FDR thresholded confidence map (right, blue): Mg^2+^ ion coordinated by E461, E416, H418 and 3 H_2_O molecules (top). Density of bound PETG ligand the 3.5/4.5 *σ*-thresholded cryo-EM map and in the 1% FDR confidence map (bottom). (b) TRPV1 channel (EMD 5778) with 5 *σ*-thresholded cryo-EM map (left) and 1 % FDR thresholded confidence map (right): selectivity filter formed by carbonyls of symmetry-related G643 residues. The presence of a putative ion is supported by the confidence map. (c) γ-secretase (EMD 3061) with 4*σ* and 5*σ*-thresholded cryo-EM map (left) and 1 % FDR thresholded confidence map (right): The confidence map reveals density for both acyl chains of phosphatidyl choline at a single threshold.

## 4 Discussion

In the current manuscript, we introduced FDR-based statistical thresholding of cryo-EM densities as a complementary tool for map interpretation. This approach is used successfully in other fields of image processing sciences (Genovese *et al.*, 2002). Based on a total of five near-atomic resolution EM maps from the EMDB model challenge (http://challenges.emdatabank.org), one intermediate resolution (6.8 Å) structure and three subtomogram averages in the resolution range between 90 and 23 Å, we showed that using 1% FDR confidence maps are well suited for detailed molecular feature detection and result in better confidence in particular for assignment of weak structural features. Although for all maps different *σ*-levels ranging between 1 and 5 could be used for the interpretation of relevant cryo-EM map features, confidence maps thresholded at a common 1 % FDR level show consistent interpretability of molecular features for these maps. The advantage of confidence maps is that they effectively separate signal from a background noise estimate by assigning a confidence scale from 0 to 1 and at 1 % FDR. This way they show consistent inclusion of signal while minimizing noise. In contrast, for cryo-EM densities small changes of the isosurface *σ*-threshold can have severe consequences for the interpretability of molecular features and bear the risk of mistakenly including noise. Therefore, confidence maps and associated FDR thresholds provide a common and conservative thresholding criterion for the interpretation of cryo-EM maps.

Included in the algorithm is a direct assessment of the signal significance with respect to background noise associated with particular density features visible in cryo-EM maps, which adds an additional objectivity to reporting of ambiguous density features. Based on these properties, high-resolution confidence maps will be helpful in initial atomic model building when no or little atomic reference structures are available and for assessment of critical details such as side chain conformations and non-protein molecules in the density. The use of these maps will improve the quality of initial atomic models before launching real-space or reciprocal atomic coordinate refinement (Murshudov, 2016; Adams *et al.*, 2010), which should proceed with sharpened or alternatively model-based sharpened maps as refinement targets (Jakobi *et al.*, 2017). The molecular interpretation based on confidence maps is not limited to maps of close-to-atomic resolution as we demonstrated its benefit for cases of intermediate resolution single-particle and subtomogram averaging with three maps ranging in resolution from 7 – 90 Å. In these cases, the interpretation of an unassigned density using a confidence level is a beneficial property in particular in the absence of atomic model information.

We also showed that the generation of confidence maps is a robust procedure. From the sharpened cryo-EM density, we compute the CDF from the solvent background, which in most cases can be approximated by a Gaussian distribution. In addition, we assume protein density to be positive as the overwhelming majority of determined atoms density resides in positive density. Moreover, we find that the region selected for noise estimation is critical as it has to contain pure noise devoid of signal. We found this particularly important for generating confidence maps from subtomogram averages with particle boundaries less well defined. Generally, when estimating background noise outside the particle, we tend to overestimate noise due to smaller ice thickness in particle regions. Smaller deviations from noise estimation show little effect on the conversion to confidence maps (**Fig. S6b**). We show that when suboptimally sharpened input maps are used to generate confidence maps, the operator avoids the common risk of mistakenly interpreting noise as signal in over-sharpened cryo-EM densities. In contrast, confidence maps generated from over-sharpened input maps will only result in insufficient declaration of density signal, which is an important safety feature. Once noise is estimated, the procedure of generating confidence maps is statistically clearly defined (Benjamini & Hochberg, 1995; Benjamini & Yekutieli, 2001) and does not contain any free parameters to optimize. Only in cases of substantial resolution variation due to molecular flexibility and computational errors, it may be required to locally adjust SNRs by including prior information through local resolution filtering. More sophisticated approaches such as amplitude scaling can also be used in cases where atomic reference structures are available. Adjusting FDR control based on prior information is routinely implemented in other applications of statistical hypothesis testing (Chong *et al.*, 2015; Ploner *et al.*, 2006). With the manuscript, we provide a program that requires a 3D volume as input and allows specification of the location of density windows used for noise estimation. The presented implementation including local resolution filtration is computationally fast, taking from 30 s to 2 min on a Xeon Intel CPU for the maps produced in this manuscript.

We presented several cases in our simulation and EMDB maps where confidence maps displayed weak structural features more clearly while minimizing the occurrence false positive pixels (**Figures 1–5**). This is a particularly useful property of confidence maps. Weak densities close to inherent noise levels are present in most cryo-EM maps and they result as a consequence of the molecular specimen as well as from the applied computational procedures. For example, they can originate from side chain mobility in the form of multiple rotamers or side-chain specific radiation damage (Fromm *et al.*, 2015; Allegretti *et al.*, 2014; Bartesaghi *et al.*, 2014). In addition, ligands including small organic compounds or larger protein complex components may have lower occupancy or partial flexibility { Zhao:2017 hi}. In many complexes, peripheral loops exposed to the solvent tend to have larger molecular flexibility than the core of the protein (Hoffmann *et al.*, 2015). We showed that thresholding confidence maps yield higher voxel detection rates than thresholding in common cryo-EM densities. We believe that is a result of the fact that the human operator prefers to recommend a more conservative *σ*-threshold to avoid excessive inclusion of noise while as a consequence one misses out on signal. Using confidence maps, this type of noise can be suppressed and as a result more reliable signal can be interpreted.

With the increasing number of near-atomic resolution cryo-EM structures, the process of building atomic models has become increasingly important but remains time-consuming and labor-intense. Confidence maps can assist the user throughout this process. In X-ray crystallography, multiple complementary maps are being used routinely in the process of model building. Real-space model building and optimization is typically performed using maximum likelihood-weighted 2mF_o_–DF_c_, assisted by mF_o_–DF_c_ difference map used to highlight errors in the model. Various forms of omit maps computed from phases of models in which a selection of atoms (e.g. a ligand) has been omitted are used to confirm the presence of ligand and ambiguous density. Similarly, confidence maps display a complementary aspect of cryo-EM maps in helping to reduce ambiguity in density interpretation of e.g. weakly bound ligands, alternative side-chain rotamers, conformationally heterogeneous structures including incomplete or flexible parts of the complex. It is evident that confidence maps would not be suitable for model refinement, as they do not discriminate the scattering mass of different atoms or relative uncertainties of atomic positions. These properties are usually modelled by atomic electron form factors and atomic displacement factors (atomic B-factors). However, owing to the increased precision of density peaks and noise suppression, it is perceivable that confidence maps could be used to guide positional coordinate refinement if implemented as a peak searching procedure. In addition, defined confidence values for density stretches should also be useful and potentially beneficial for automated model building approaches. Interpreting cryo-EM densities by means of an atomic model is often the final step of a cryo-EM experiment. In practice, atomic models are even used as a validation tool to examine density features for side chains at expected positions. One of the key advantages of the here proposed confidence maps is that they can be generated without prior knowledge of an atomic model. As the conversion of cryo-EM densities to FDR controlled maps is a conceptually simple and computationally straightforward, confidence maps could be routinely consulted for providing complementary information of statistical significance during the intricate process of interpreting ambiguous densities in cryo-EM structures resulting from molecular flexibility or partial occupancy.

## Author contributions

M.B. and C.S. initiated the project. M.B. developed and implemented the code for the algorithm. A.J.J. helped with structure comparison and implementation including LocScale integration. C.S. supervised the project. M.B. and C.S. wrote the manuscript with input from A.J.J.

## Acknowledgements

M.B. has been supported by the EMBL International PhD Programme. We thank Martin Beck (EMBL) for critical reading of the manuscript and the thesis advisory committee members Wolfgang Huber and Rob Russel (Heidelberg University) for stimulating discussions. We are grateful to Thomas Hoffmann and Jurij Pečar (IT Services) for set-up and maintenance of the high-performance computational environment at EMBL.

## 5 Appendix

### 5.1. Statistical model

For each voxel in the reconstructed 3D volume, where the voxels are indexed with *i,j,k*, the intensity *X _i,j,k_* is modeled as

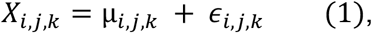

with ϵ_*i,j,k*_ a real valued random variable representing the background noise with mean *μ*_0,*i,j,k*_ ∈ ℝ and variance 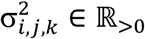 and where *μ_i,j,k_* ∈ ℝ is the true intensity as observed without background noise.

We developed an algorithm by means of multiple hypothesis testing, that controls the maximum amount of false positive signal in the map, i.e. the FDR with respect to background noise. First, we limit the tested voxels to the reconstruction sphere and voxels located outside of a diameter larger than the box size are disregarded as they arise from a smaller subset of averaged images than the voxels inside. Second, we focus on the detection of voxels with positive deviations from background noise (see section 3.2). In addition, voxels that contain significant signal are affected by further sources of noise like flexibility, incomplete binding of ligands and structural heterogeneity, leading to intensity variations of the signal. Consequently, these sources lead to an increase of the variance for these voxels as part of incoherent signal, which we do not consider here as it is going beyond the scope of detecting signal beyond background. Background noise of experimental cryo-EM data, however, poses principal challenges to the statistician, as it can result in non-uniform distributions across the map: although background noise variances from images of uniform noise over the pixels can be assumed uniform over the central sphere (**Fig. S3c right**), background noise outside the particle is higher when compared with background noise affecting the particle itself due to solvent displacement and variations of relative ice thickness at the particle (Penczek *et al.*, 2006). Therefore, estimating noise in the solvent region outside the particle could lead to an overestimation of the actual influence of the background noise on the particle (see section 3.4). Although this may cause several problems for comprehensive probabilistic modelling, these estimates can be interpreted as conservative bounds for the signal significance of the particle over background noise. For this reason, we use multiple hypothesis testing in order to calculate these upper bounds for detection errors of false positive rates, as we prove in Proposition 1. In cases when alternative noise estimates are available, they can be supplied as additional input to the procedure in order to generate confidence maps.

For each voxel a *z*-test is carried out, which identifies significant deviations from background noise. The value of the test statistic *Z* at each voxel is then given as

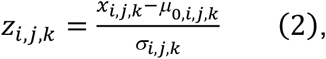

where *x_i,j,k_* ∈ ℝ is the reconstructed mean intensity at the respective voxel. We are testing for true intensity *μ_i,j,k_* higher than 0, thus the null and alternative hypotheses for each voxel become

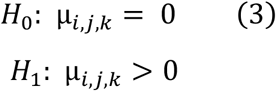

The null hypothesis *H*_0_ states that the true intensity *μ_i,j,k_* at the respective voxel is 0, i.e. no signal beyond background noise, while the second hypothesis *H*_1_ states the deviation towards higher values. Testing for deviations towards negative values, i.e. negative densities, is easily accomplished in this setting by multiplying the normalized map intensities *z_i,j,k_* with –1, leading to a left-sided test procedure. Both options can be chosen by the user.

Under the null hypothesis *H*_0_ and by approximating the background noise with a Gaussian distribution (Kucukelbir *et al.*, 2014; Vilas *et al.*, 2018), the test statistic *Z* follows a standard Gaussian distribution. The *p*-values in our procedure are then calculated as

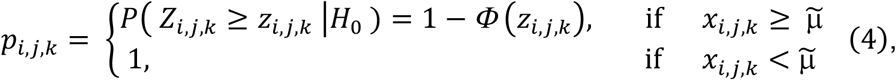

with *Z_i,j,k_* being the random variable representing the test statistic at voxel *i,j,k*, *z_i,j,k_* the particular realization, *μ̃* the background noise as estimated from the solvent area and the cumulative distribution function Φ() of the standard Gaussian distribution. Alternatively, *p*-values can also be calculated in a non-parametric way without any assumptions about the underlying background noise distribution by simply replacing the cumulative distribution function Φ() of the standard Gaussian distribution with the empirical cumulative distribution function *F*() estimated from the sample of background noise, given as

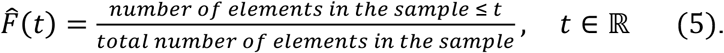

This allows the complete procedure to be carried out without any distribution assumptions. However, comparisons show that the background noise can be well approximated with a Gaussian distribution even in the tail areas, which are most important for the calculation *p*-values (see section 3.2, **Fig. 1b and S3a**). The respective method for *p*-value calculation, i.e. non-parametric or with Gaussian assumption, can be chosen by the user. All presented cases in the manuscript, if not stated differently, were calculated with the assumption of Gaussian distributed background noise. Note, the here defined *p*-values differ only marginally from the *p*-values commonly used for one-sided testing in a way that for all voxels with intensities smaller than the expected mean noise level *μ*_0_ their value is here set to one. This definition allows the control of the FDR in the more general setting of allowed overestimated mean and variance (see Proposition 1).

### 5.2. Multiple testing correction

The respective hypothesis tests are applied to each voxel in the 3D volume. To account for the multiple testing problem with up to more than a million tests, we choose to control the FDR. Control in this context is meant in giving upper bounds for the occurring error. The FDR is defined as the expected amount of false rejections, i.e.

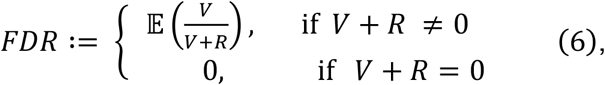

with *V* ∈ ℕ. the number of false rejections, *R* ∈ ℕ. the number of true rejections and 𝔼 () denotes the expectation value. Due to dependencies between hypotheses at voxels close to each other, we choose the Benjamini-Yekutieli procedure (Benjamini & Yekutieli, 2001), giving an FDR-adjusted *p*-value for each voxel, which are often referred to as *q*-values. To describe the adjustment of *p*-values according to Benjamini and Yekutieli in more detail and for the ease of notation, we will now use a sequence of voxels from the map and denote the number of hypotheses, i.e. tested voxels, with *m*. The *p*-values *p_i_*, *i* = 1, …, *m* are then sorted, from small to large, resulting in sorted *p*-values *p*_(*i*)_, *i* = 1, …, *m*. *q*-values are then calculated as

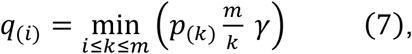

with *m* the number of hypotheses, *k* a running index and 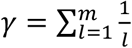 By recognizing the correct index in the sequence of voxels for each index (*i*), *i* = 1, …, *m* in the sorted array and subsequent conversion into the 3D volume, we can assign each voxel position *i,j,k* its corresponding *q*-value. In order to interpret the resulting map, the *q*-value for each voxel then gives the minimal FDR that has to be imposed at the thresholding in order to call the respective voxel a significant deviation from the background. The final value associated with voxel 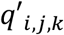, is then calculated as

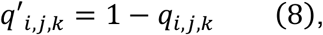

where *q _i,j,k_* is the *q-*value at the voxel indexed with *i,j,k.* Thus, visualization of the map at a value of 0.99 corresponds to a maximal FDR of 1%, or a minimal PPV of 99%, and therefore means that from all the visible voxels at this threshold, a maximum of 1% are expected to be background noise.

Next, we show that the presented procedure with *p*-values as defined above controls the FDR even in the case of overestimated background noise, i.e. by using the possibly overestimated background noise estimates from the solvent area in Equation (2) for all voxels.

#### Proposition 1

*Consider Gaussian distributed random variables representing the background noise at all voxels i,j,k in the 3D map with true mean μ_0,I,j,k_* ∈ ℝ *and variance* 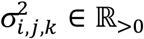*. Moreover, let* μ̃ ≥ μ_0,i,j,i_*and* 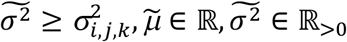. *for all i,j,k, the overestimated background noise parameters. Then* 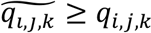*, where* 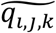 *corresponds to the q-value as defined in Equation (7) and calculated with our procedure with parameters* 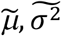*, and q*_,*i,j,k*_ *the q-value, as obtained with the true parameters μ*_0,*i,j,k*_ and 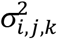.

<u>Proof</u>:

In order to prove the statement, we will now recapitulate the algorithm and prove the inequality at all necessary steps. We start showing that the true *p*-value at voxel position *i,j,k*, *p_i,j,k_*, is smaller when compared with the *p-*value 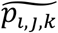 calculated from the overestimated background noise parameters using Equation (4). In other words, we want to show that 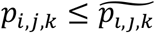 or equivalent to that, 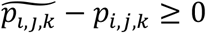. If *x_i,j,k_*<*μ̃* then the statement is trivial, because 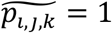 and *p_i,j,k_* ≤ 1, which is a general property of *p-*values.

For *x_i,j,k_*≥*μ̃*, considering Equations (2) and (4), it follows:

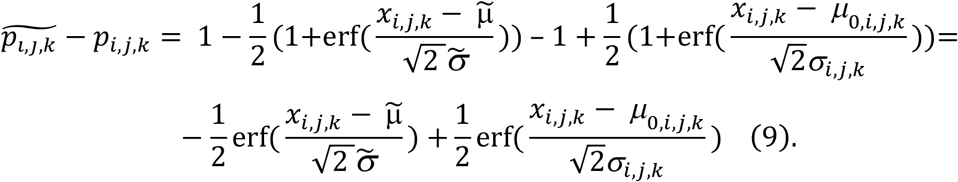

As the error function erf() is monotonically increasing, it is sufficient to show that 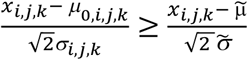. Because *x_i,j,k_*−*μ̃*≥0 and thus also, *x_i,j,k_*−*μ*_0,*i,j,k*_≥0 as well a *σ̃*≥*σ_i,j,k_*,we have

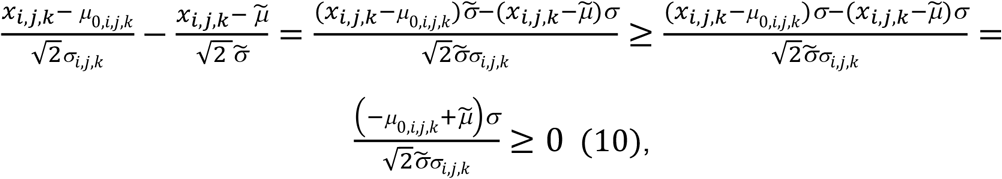

where in the last inequality it was used that *μ̃*≥*μ*_0_,_*i,j,k*_ and σ̃≥σ_*i,j,k*_> 0. This gives the desired result of 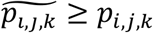.

Recapitulating the calculation of *q*-values in Equation (7) together with the conversion of the 3D volume to a sequence, it follows:

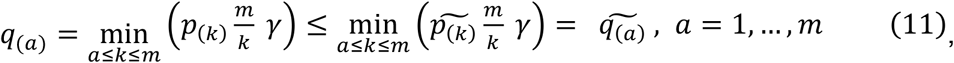

with *m* the number of hypotheses, *k* a running index and 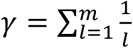. This gives the desired result:

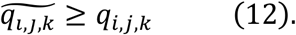

As the Benjamini-Yekutieli procedure controls the FDR when using true parameters, our procedure (i.e. Benjamini-Yekutieli applied to the modified *p*-values) will give a more conservative estimate of the FDR (as shown in Proposition 1). Therefore, our algorithm controls the FDR sufficiently well by giving an upper conservative bound for the FDR. Thus, Propostion 1 states that even in the setting of non-uniform background noise with higher noise levels in the region of background noise estimation, the FDR is controlled and thus robust in the sense that the maximum FDR is still guaranteed. Furthermore, it has to be mentioned that estimates of the background noise levels are not the only factor contributing to FDR estimation. Both the number voxels as well as their dependencies within the map have an important influence and are considered in the FDR-adjustment. This makes the generation of confidence maps even with severely overestimated background noise parameters a powerful procedure (**Fig. S6**), where powerful is used here in its statistical sense of decreasing the error of missing true signal. However, the power of the procedure can be even increased, i.e. the amount of true missed signal reduced while controlling the FDR, by including information about local resolutions, cutoffs in reciprocal space where no signal is expected beyond, while, at the same time, controlling the FDR.

### 5.3. Testing with local filtering

In the presence of extreme resolution variation, using uniformly sharpened and filtered maps will lead to confidence maps of insufficient representation of features in both areas of either lower than the average B-factor or higher than the average B-factor.

Therefore, in the next two sections, we will show how noise levels can be locally adjusted and subsequently estimated by inclusion of local resolution information as well as atomic B-factors and how this can be used to increase the power to detect weaker features while controlling the FDR. Local filtration of EM maps according to the local resolution (Cardone *et al.*, 2013) has been shown to be a powerful approach as it leads to local reductions of background noise. These variations of noise levels between different voxels at different resolutions from local filtering, can be also accounted for in the generation of confidence maps. For each voxel, a map duplicate volume is filtered at the corresponding resolution and the noise distribution estimated from the solvent area outside the particle. This procedure results in three 3D maps, the estimates of local variances of the background noise at each voxel after local filtration, the estimates of local means of the background noise at each voxel after local filtration and the locally filtered map. These three maps are subsequently used for the testing procedure. Thus, the value of the test statistic (2) is calculated by

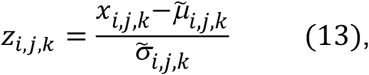

where *x_i,j,k_* ∈ ℝ is the intensity of the locally filtered map at voxel *i*, *j*, *k* and *μ̃_i_*_, *j*, *k*_ ∈ ℝ and *σ̃_i_*_, *j*, *k*_ ∈ ℝ_>2._ are the local mean and standard deviation estimate of the background noise at the respective voxel. All subsequent steps of the algorithm remain identical as well as the validity of Proposition 1.

### 5.4. Testing with local amplitude scaling

As for the local filtration, local amplitude scaling gives rise to varying noise levels at different voxels. In order to obtain both mean and variance estimates for each voxel after local amplitude scaling, a duplicate window outside the particle containing pure noise is scaled according to the rolling window used in local amplitude scaling for each voxel, i.e. the amplitudes of the Fourier transform of the box containing pure noise at frequency *s*, denoted as *F*_noise_(*s*), are multiplied with a frequency dependent sharpening factor *k*(*s*) ∈ ℝ_≥0_, which is consequently given as*k*(*s*)

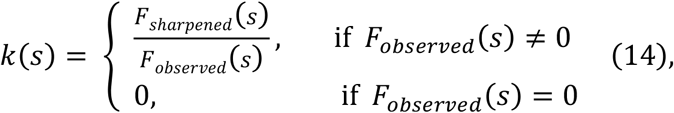

where *F_sharpened_*(*s*) ∈ ℝ_≥0._ and *F_observed_*(*s*) ∈ ℝ_≥0._ are rotationally averaged amplitudes of the Fourier transform at frequency *s* given at the respective rolling window for the sharpened and the observed experimental map, respectively. The noise distribution is then estimated from the scaled noise sample. In analogy to the case of locally filtered maps, this procedure results again in three 3D maps of estimated means, variances and intensities of the locally sharpened map for each voxel that can be incorporated with Equation (13) in the testing procedure. Proposition 1 remains valid.

---

**Fig. S1.**
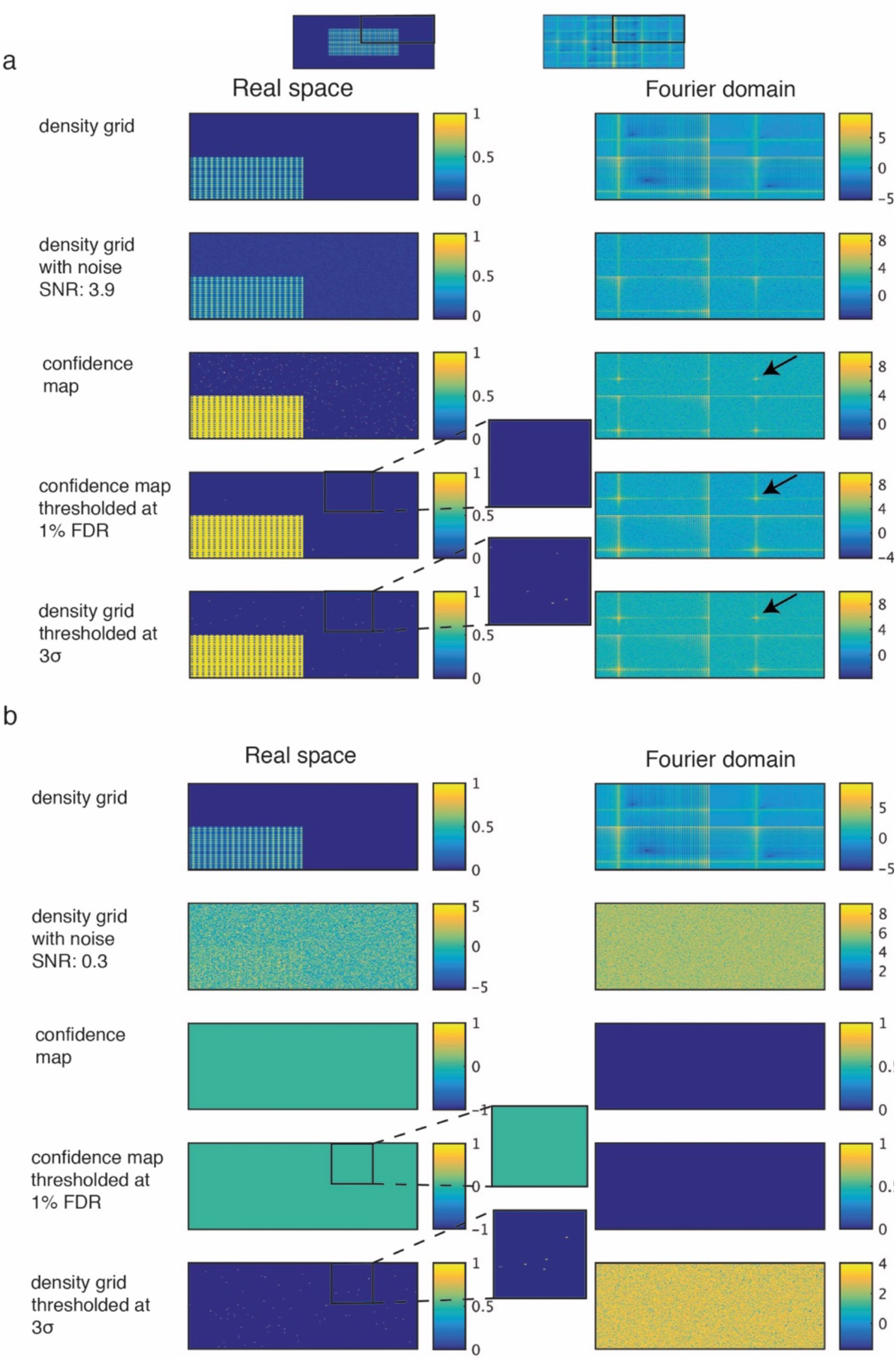
Comparison of *σ* and FDR thresholding of simulated density grids with varying signal-to-noise ratios. Thresholding with simulated density grids at signal-to-noise ratios and variance of (a) 3.9 (0.01) and (b) 0.3 (1.33), respectively. The same simulations as in Fig. 1c are repeated with lower and higher variances of the background noise. At low signal-to-noise ratio, the 1% FDR thresholding is devoid of false positives whereas conventional 3*σ*-thresholding approach yields many false positive pixels (zoomed inset).

**Fig. S2.**
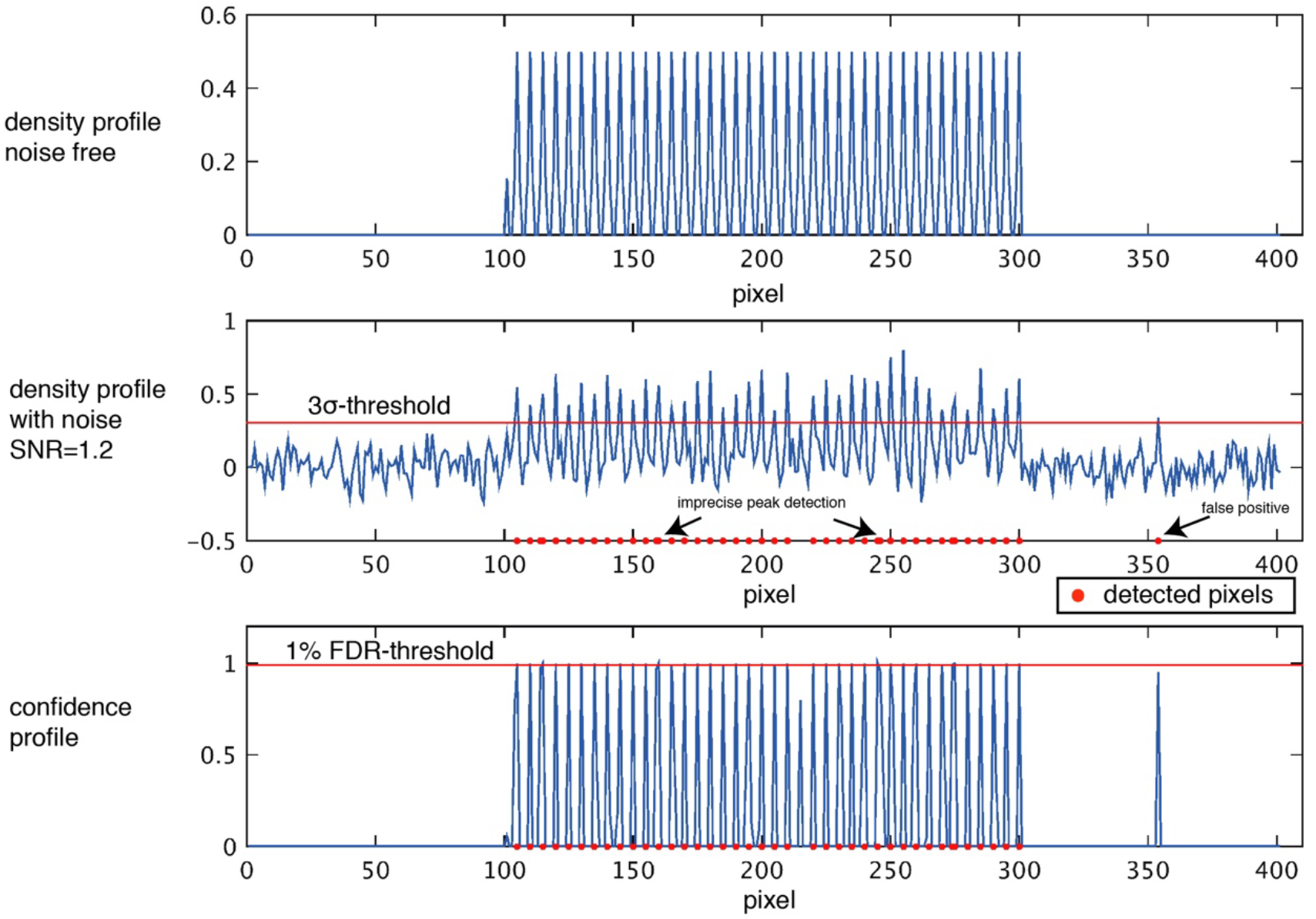
Effect of *σ* and FDR thresholding on 1D density profiles. One-dimensional stacked plots of grid density with noise-free original (top), at signal-to-noise ratio of 1.2 (center) and confidence map (bottom). The noisy density grid is thresholded at 3*σ* and the confidence map is thresholded at 1 % FDR. Conventional 3*σ*-thresholding yields higher rates of false positives and some imprecise peak positions (arrows).

**Fig. S3.**
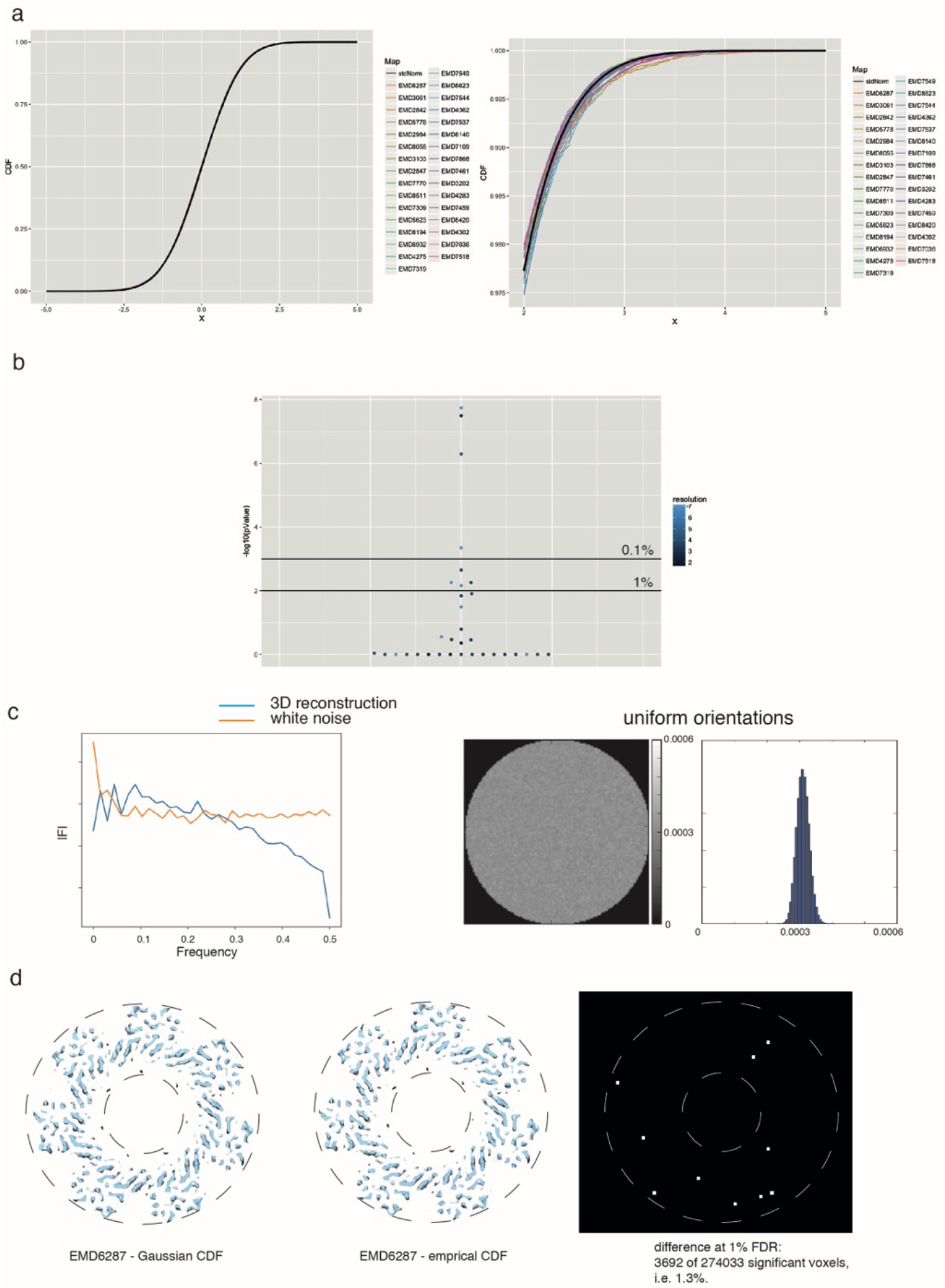
Analysis of normality of cryo-EM densities. (a) Left. Overlay of 32 cumulative density functions (CDF) derived from the above EMDB entries with ideal Gaussian CDF in black. Right. Zoomed inset to better highlight small differences. (b) 32 map entries are assessed with respect to normality according to the Anderson-Darling test, significance thresholds are displayed 1.0 and 0.1 % respectively. (c) Left. Rotational power spectrum of a 3D reconstruction of white noise images in comparison with pure white noise spectrum. Right. Slice through 3D volume of variances estimated from 900 independent reconstructions from Gaussian white noise images with similar uniform orientations together with a histogram of the estimated variances, showing that background noise can be assumed uniform over the central sphere in the reconstructed volume. (d) Cross-sectional view of confidence maps generated of EMD6287 using Gaussian and empirical CDF. Difference map between 1 % FDR binarized confidence maps in the respective image slice.

**Fig. S4.**
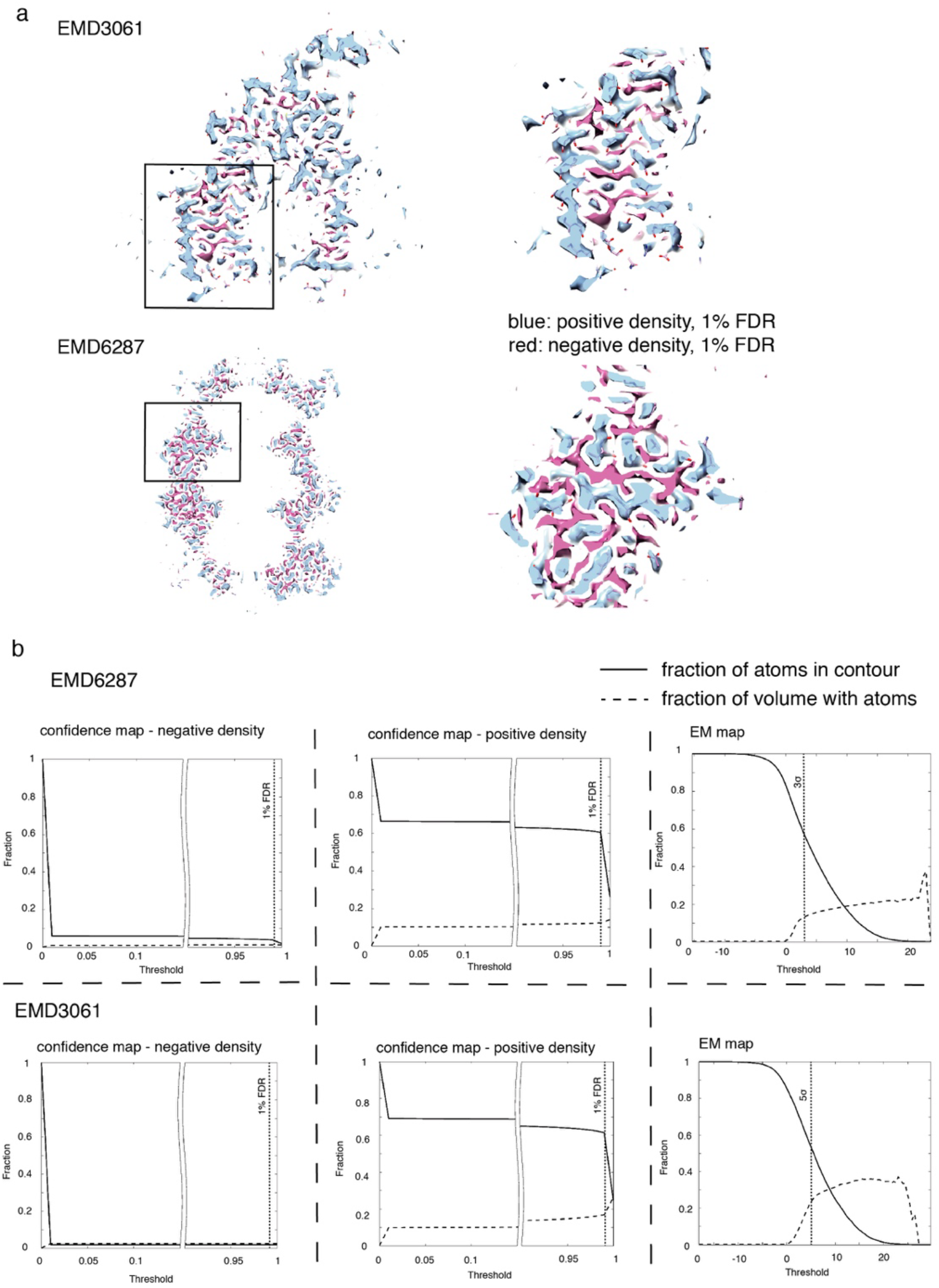
Analysis of positive and negative densities using confidence maps. (a) Overlay of 1% FDR positive (blue) and negative (red) confidence maps from original and inverted densities of EMD3061 (top) and EMD6287 (bottom) respectively. (b) Comparison of detected signal with corresponding atomic models by determining the fraction of overlap of atoms with volume and fraction of volume with atoms as a function of threshold for negative (left), positive (center) confidence maps and cryo-EM maps (right), respectively.

**Fig. S5.**
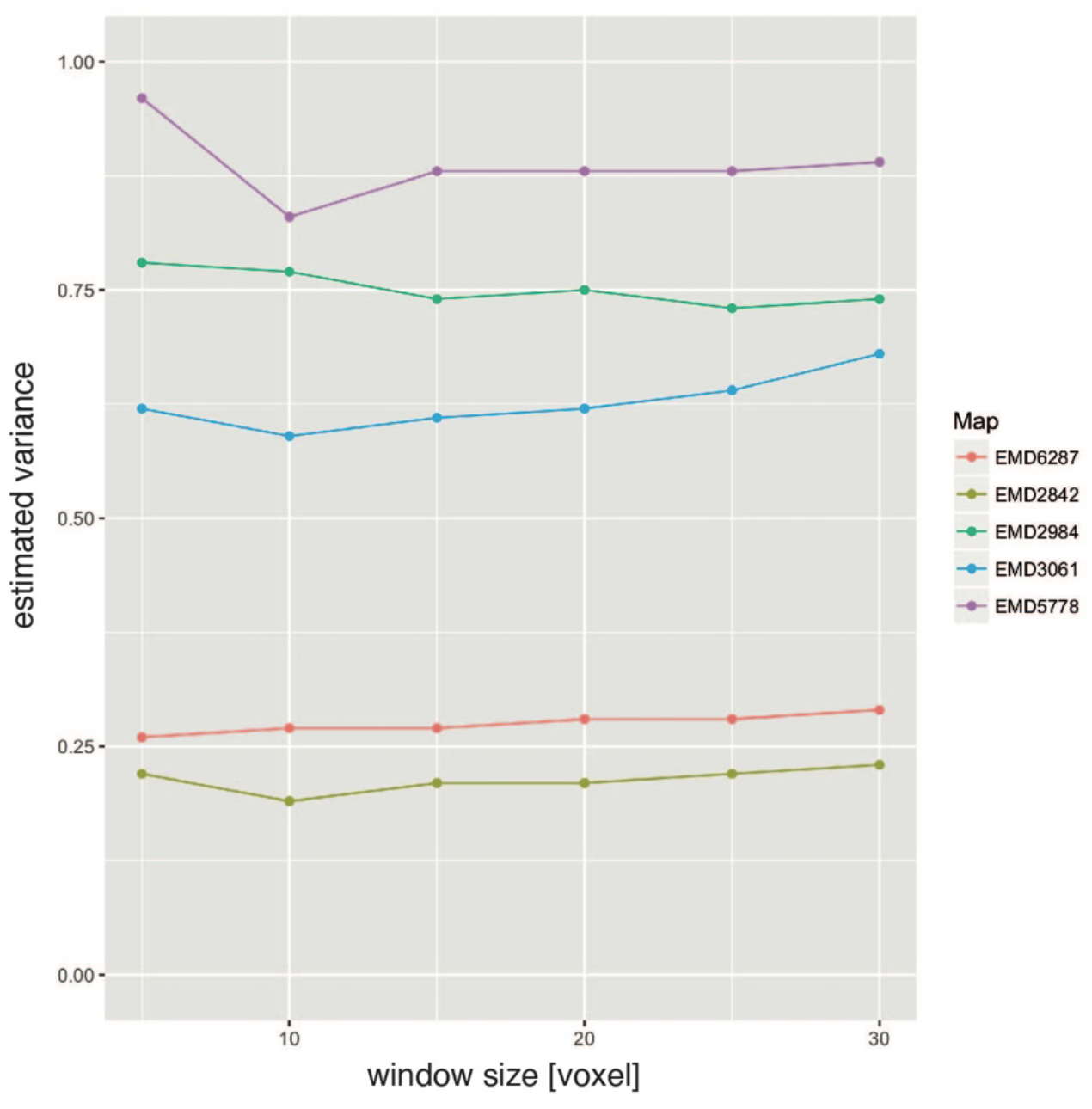
Effect of window size on estimated variance. Estimated variance is stable with increasing window size from 5 to 30 voxels for a series of EMD entries.

**Fig. S6.**
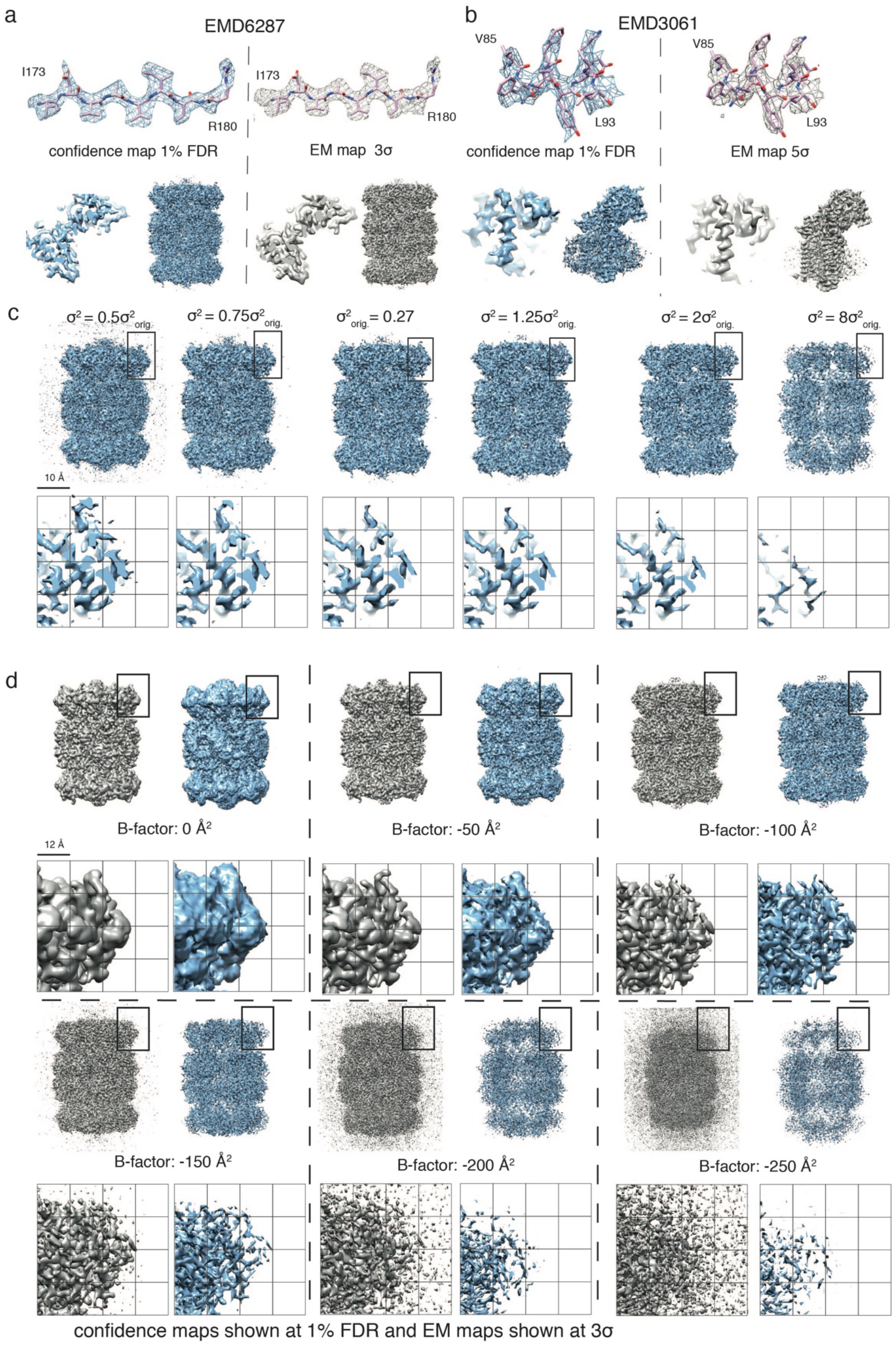
Confidence maps and effect of incorrect noise estimation. (a) 20S proteasome map (EMD 6287) comparison of 1% FDR density (left) and 3*σ*-thresholded map (right). Shown are molecular details from I173 – R180 (top), slice view (bottom left) and side view (bottom right) of density. (b) γ-secretase map (EMD 3061) comparison of 1% FDR confidence map and 5*σ*-thresholded map. (c) Six confidence maps of 20S proteasome (EMD-8267) including magnified inset based on incorrect variance estimation: 1^st^ and 2^nd^ left noise is underestimated by 0.5 and 0.75 times the variance (*σ*^2^). In comparison with the correctly estimated noise (3^rd^), they show excessive noise features declared as signal at 1 % FDR. When noise is overestimated, which is more likely for cryo-EM maps, confidence maps are quite insensitive to changes in map appearance. For multiples like 1.25*σ*^2^ and 2*σ*^2^ no apparent density changes become visible (4^th^ and 5^th^) unless strong overestimation like 8*σ*^2^ (6^th^) leads to disappearance of map features at a 1 % FDR threshold. (d) When applying a series of B-factors to the 3D reconstruction of the 20S proteasome map, we see that with higher B-factors, sharpened EM densities become dominated by noise whereas corresponding confidence maps displayed at 1 % FDR show disappearance of significant features thereby avoids over-interpreting noise features.

**Fig. S7.**
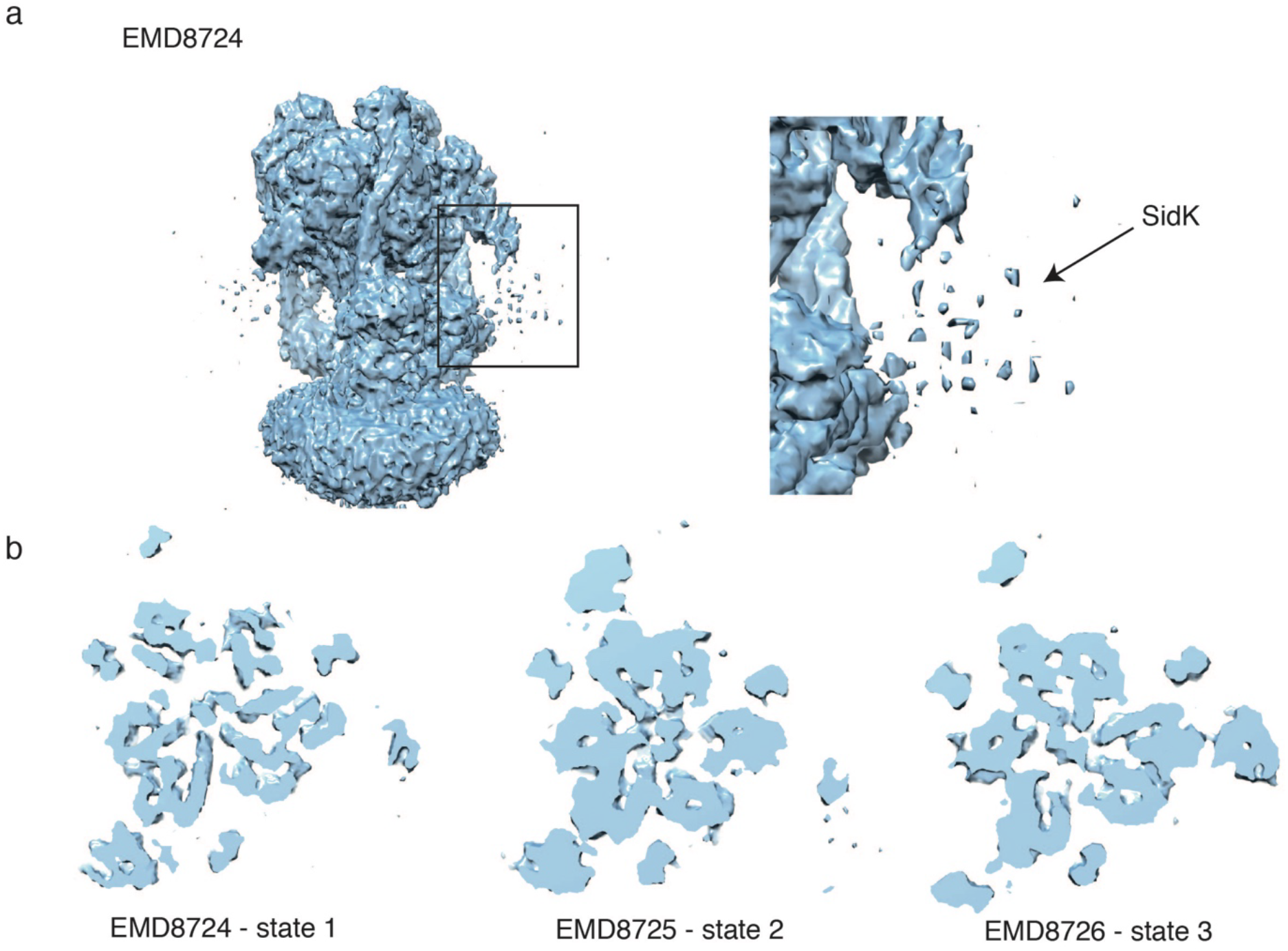
Confidence maps of compositionally and conformationally heterogeneous complexes. (a) Confidence map of yeast V-ATPase with Legionella pneumophila effector SidK (EMD8724) at 1% FDR (left) together with a zoom on the flexible domains of SidK (right). The confidence map shows significant density for the flexible domains, however, not as continuous density. (b) Slices through confidence maps of 3D classified cryo-EM maps. Different rotational states can be resolved in the confidence maps.

**Fig. S8.**
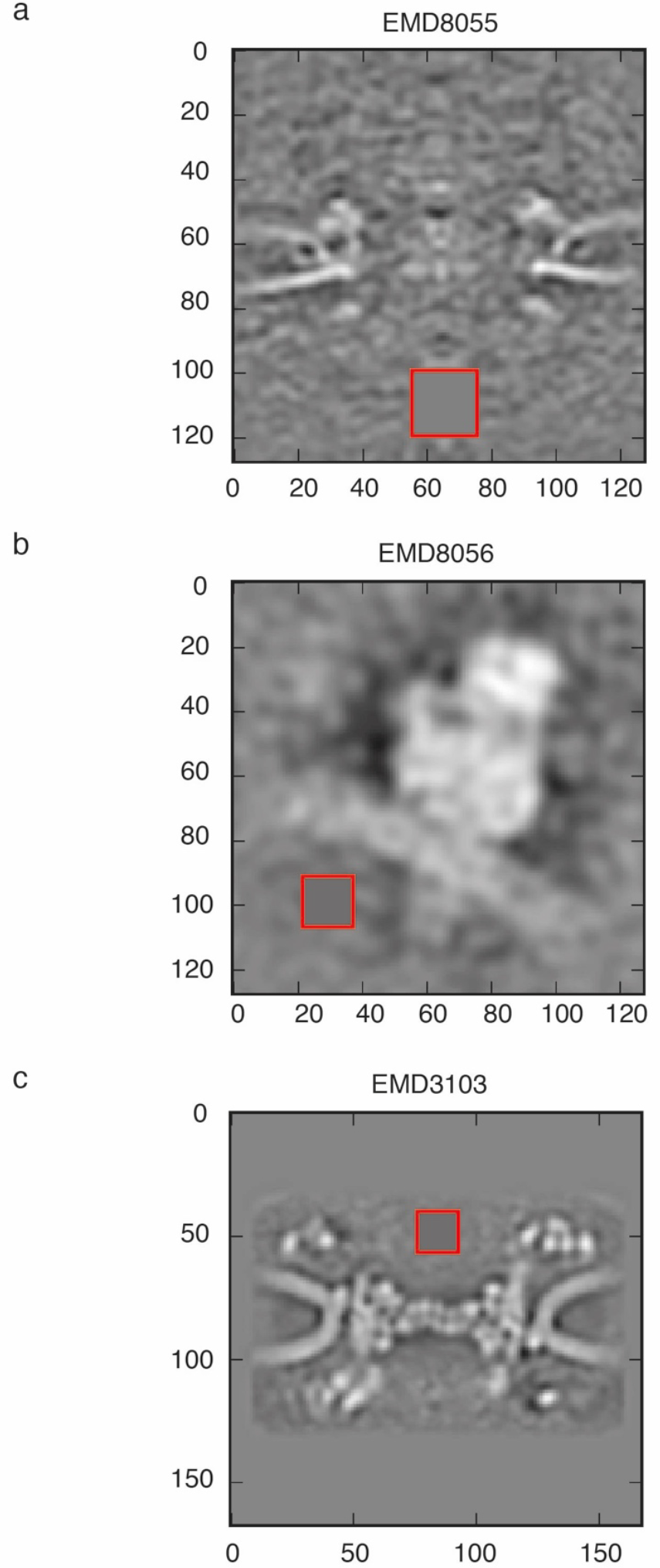
Noise estimation in subtomogram averages. Gray-scale density slices with red windows for the voxel region used for variance estimation: (a) EMD 8055: nuclear pore from HeLa cells by FIB-SEM, (b) EMD 8056: ER-associated ribosomes, (c) EMD3103: 23 Å resolution nuclear pore subtomogram average.

**Fig. S9.**
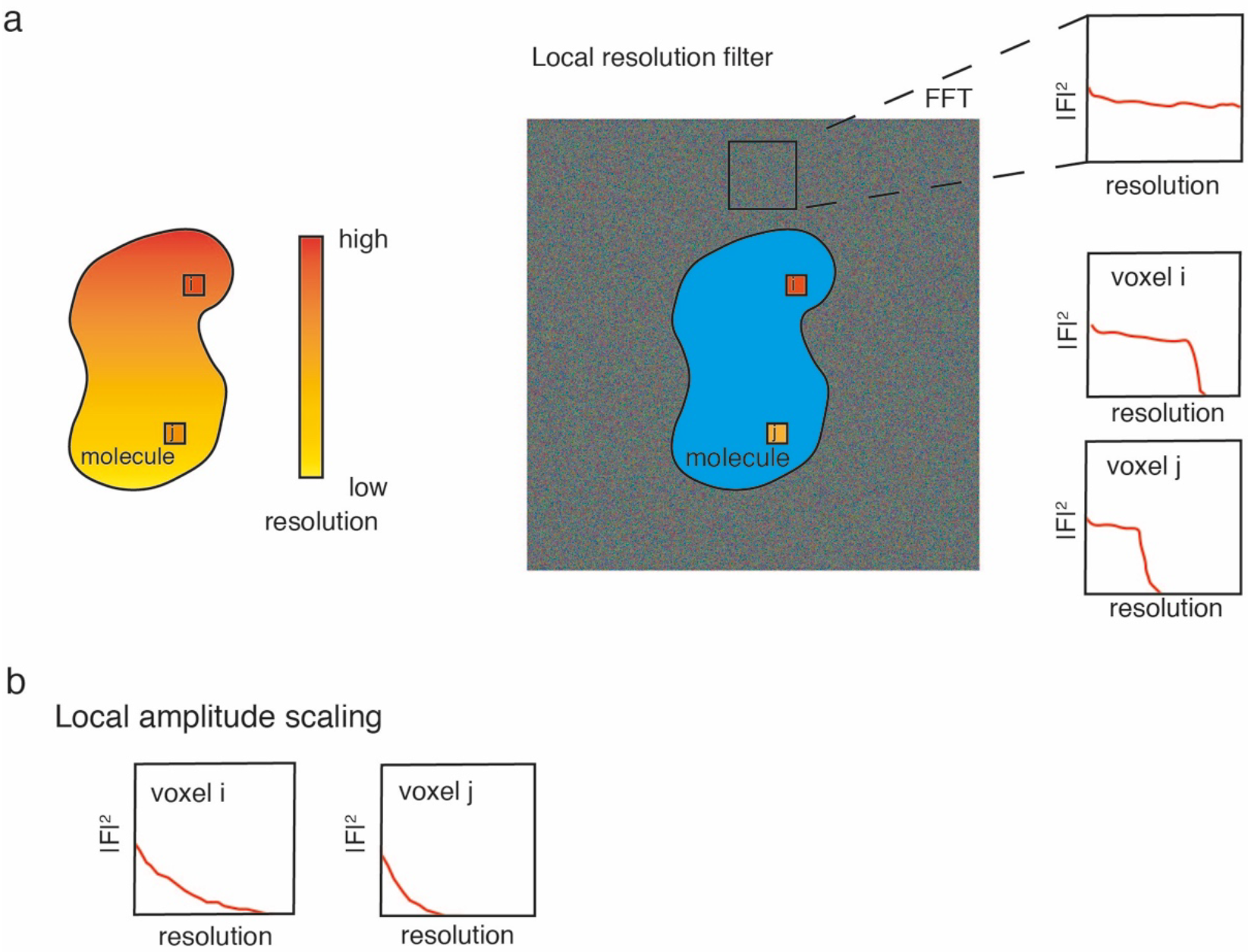
Variance adjustment based on local resolution and local amplitude profile. (a) Adjusting the local signal-to-noise ratio based on local resolution measurements: for each voxel, the background windows are filtered according to the local resolution at the respective voxels in order to estimate the noise levels of each voxel in the locally filtered map. (b) In analogy, local sharpening is applied to background noise in order to estimate resulting local noise distributions.

**Fig. S10.**
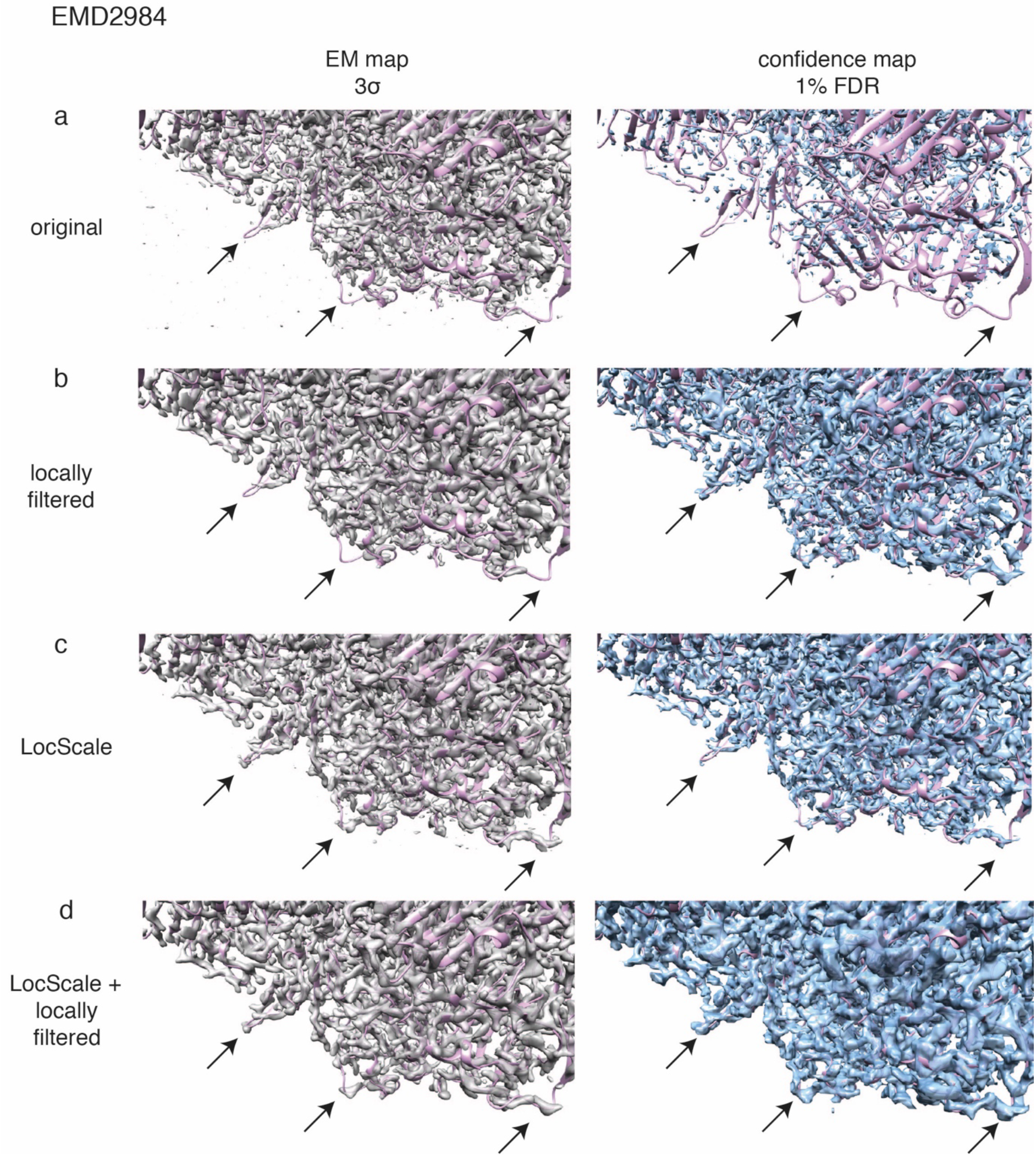
Effect of local variance adjustments on confidence maps. β-galactosidase (EMD 2984) cryo-EM map at 3.0*σ* threshold (left, gray) and 1 % FDR confidence map based on different post-processing methods (right, blue). Global sharpening with uniform filtering, local filtering based on local resolution measurements, local sharpening and the combination of local sharpening with local filtering were compared. Confidence maps were generated with local noise estimate based on local resolution measurement, locally scaled window from a model reference structure and the combination of both, which in this case shows the best preservation of molecular density with respect to confidence.

